# Mitophagy antagonism by Zika virus reveals Ajuba as a regulator of PINK1-Parkin signaling, PKR-dependent inflammation, and viral invasion of tissues

**DOI:** 10.1101/2021.01.29.428870

**Authors:** Sanket S. Ponia, Shelly J. Robertson, Kristin L. McNally, Gail L. Sturdevant, Matthew Lewis, Forrest Jessop, Catherine M. Bosio, Catherine Kendall, Dylan Gallegos, Arielle Hay, Cindi Schwartz, Rebecca Rosenke, Greg Saturday, Craig Martens, Sonja M. Best

**Affiliations:** Innate Immunity and Pathogenesis Section, Laboratory of Virology, Rocky Mountain Laboratories, National Institute of Allergy and Infectious Diseases, NIH, Hamilton, MT 59840; Immunity to Pulmonary Pathogens Section, Laboratory of Bacteriology, Rocky Mountain Laboratories, National Institute of Allergy and Infectious Diseases, NIH, Hamilton, MT 59840; School of Molecular and Cellular Biology, University of Leeds, Leeds, United Kingdom; Research Technology Branch, Rocky Mountain Laboratories, National Institute of Allergy and Infectious Diseases, NIH, Hamilton, MT 59840; Rocky Mountain Veterinary Branch, Rocky Mountain Laboratories, National Institute of Allergy and Infectious Diseases, NIH, Hamilton, MT 59840

## Abstract

Dysregulated inflammation dominated by chemokine expression is a key feature of disease following infection with the globally important human pathogens, Zika virus (ZIKV) and dengue virus, but a mechanistic understanding of how pro-inflammatory responses are initiated is lacking. Mitophagy is a quality control mechanism that regulates innate immune signaling and cytokine production through selective degradation of damaged mitochondria. Here, we demonstrate that ZIKV NS5 antagonizes mitophagy by binding to the host protein Ajuba and preventing its translocation to depolarized mitochondria where it is required for PINK1 activation and downstream signaling. Consequent mitophagy suppression amplified the production of pro-inflammatory chemokines through PKR sensing of mitochondrial RNA. In Ajuba^−/−^ mice, ZIKV induced early expression of pro-inflammatory chemokines associated with significantly enhanced dissemination to tissues. This work identifies Ajuba as a critical regulator of mitophagy, and demonstrates a role for mitophagy in limiting systemic inflammation following infection by globally important human viruses.

## INTRODUCTION

Zika virus (ZIKV), a mosquito-borne flavivirus, was first isolated in Uganda (1957) but underwent an explosive emergence first affecting pacific islanders of Yap (2007) and French Polynesia (2013), before being introduced into the Americas via Brazil (2014). Although approximately 80% of infections in adults are asymptomatic or mild, infection can cause the neurological disorder Guillain-Barré syndrome or result in severe congenital neurological sequelae during pregnancy (Pierson and Diamond, 2020). Serum biomarkers of acute phase immune responses are dominated by chemokines that become highly elevated in severe disease (Foo et al., 2018; Kam et al., 2017; Michlmayr et al., 2017; Michlmayr et al., 2020; Naveca et al., 2018).

Chemokine and other pro-inflammatory responses are critical in leucocyte recruitment and control of virus infection, although uncontrolled or excessive inflammatory responses are drivers of immunopathology (Melchjorsen et al., 2003). Therefore, determining the innate immune signaling mechanisms that drive pro-inflammatory chemokine expression following flavivirus infection is key to understanding flavivirus pathogenesis and development of therapeutics.

Mitochondria are critical to the coordination of interferon (IFN) and inflammatory responses to infection with RNA viruses through two major mechanisms. The first is as a membrane platform to relay initial detection of viral double stranded RNA (dsRNA) by the RIG-I-like helicases (RLR), RIG-I and Mda5. Downstream signal transduction requires mitochondrial antiviral signaling protein (MAVS) on the surface of mitochondria to coordinate the transcriptional activation of type I and III IFNs (Mills et al., 2017). The second role of mitochondria is through release of danger associated molecular patterns (DAMPs) (West et al., 2015) including mitochondrial DNA, RNA, and cardiolipin. These signal through various pattern-recognition receptors (PRRs) including cGAS-STING, Mda5, protein kinase R (PKR), and the inflammasome (Youle, 2019). Both RLR- and DAMP-dependent responses are regulated by dynamic remodeling of the mitochondrial network including fusion to form elongated networks, fission to fragment mitochondria, and mitophagy to selectively remove irreparably damaged mitochondria through autophagolysosomal degradation (Harper et al., 2018). Failure to eliminate damaged mitochondria drives chronic inflammation that is linked to the neurodegenerative diseases such as Parkinson’s disease (PD) and Alzheimer’s disease (AD) (Mottis et al., 2019; Sliter et al., 2018).

The most well characterized pathway of mitophagy is governed by two genes, the kinase PTEN-induced putative kinase 1 (*PINK1*) and the E3 ubiquitin ligase Parkin (*PRKN*), that are mutated in familial forms of PD (Harper et al., 2018; Sekine and Youle, 2018). Following loss of mitochondrial potential, PINK1 accumulates on the surface of depolarized mitochondria where it activates itself through phosphorylation and then phosphorylates ubiquitin (Ub) on Ser65 (pS65-Ub). pS65-Ub recruits and retains Parkin at the mitochondria, enabling Parkin to be phosphorylated and activated by PINK1. Parkin then works cooperatively with PINK1 to build pS65-Ub chains on mitochondrial outer membrane proteins and recruit the autophagolysosomal machinery that ultimately results in mitochondrial clearance (reviewed in (Harper et al., 2018; Sekine and Youle, 2018)). In the context of RLR signaling, MAVS activation results in oxidative damage that triggers mitophagy as one mechanism to resolve this response (Song et al., 2020). To date, viruses have only been shown to increase mitophagy in order to dampen RLR signaling. However, while examples of viruses that inhibit mitophagy are not known, retention of damaged mitochondria in infected cells may have major implications to the host inflammatory response.

Here we reveal that ZIKV antagonizes PINK1-Parkin signaling to suppress mitophagy and that this is directly translated to pro-inflammatory chemokine expression through PKR. We show that the flavivirus nonstructural protein 5 (NS5) interacts with the cellular protein Ajuba to suppress PINK1-Parkin dependent mitophagy. Ajuba belongs to the LIM family of proteins whose demonstrated functions include the relief of Ser/Thr kinase autoinhibition and as scaffolding adaptor proteins to promote association of multi-protein complexes (Jia et al., 2020). Ajuba has been identified as an activator of mitotic kinases, including Aurora-A and CDK1 (Chen et al., 2016; Hirota et al., 2003), that also have central functions in mitochondrial dynamics (Archer, 2013). Our results demonstrate that Ajuba is recruited to mitochondria following various inducers of mitochondrial stress including MAVS activation where it is required for efficient activation of PINK1. The consequences of mitophagy antagonism to ZIKV infection include increased induction of pro-inflammatory chemokines considered to be biomarkers of ZIKV disease severity in humans. We also show that these chemokine responses are PKR-dependent, the activation of which occurs in response to increased release of mitochondrial RNA (mtRNA). Finally, we show that suppressed mitophagy results in earlier amplification of inflammatory responses in response to ZIKV infection in mice, and facilitates increased viral invasion of tissues. Together, this work identifies Ajuba as a critical regulator of mitophagy and demonstrates a systemic role of mitophagy in limiting inflammation and protection from globally important human viruses in vivo.

## RESULTS

### Ajuba negatively regulates MAVS expression dependent on mitophagy

An interaction between Ajuba and the NS5 proteins of flaviviruses was implicated from a yeast 2-hybrid performed using a cDNA library from mouse macrophages (Taylor et al., 2011) (this aspect is experimentally addressed in Figure 4B). As ZIKV NS5 is multifunctional protein with central roles in antagonism of host IFN responses (Grant et al., 2016; Xia et al., 2018), we first examined the potential of Ajuba to regulate RLR-MAVS signaling (Stone et al., 2019; Suthar et al., 2013). IFNβ expression can be induced through this pathway in tissue culture by overexpression of MAVS. *IFNB* mRNA driven by MAVS was reduced by ~84% in the presence of Ajuba (Figure 1A) and was associated with reduced MAVS expression (Figure 1B). Expression of a related LIM family member, LIMD1, had similar but less pronounced effects on expression of both MAVS and *IFNB* mRNA, suggesting that members of the LIM family may redundantly regulate RLR signaling at the level of MAVS. Replication of vesicular stomatitis virus (VSV), often used as a biological indicator of IFN sensitivity, was lower in human A549 cells depleted for *AJUBA* or *LIMD1* mRNA expression by siRNAs consistent with a role for Ajuba in negative regulation of RLR signaling (Supplemental Figure 1). We also observed recruitment of Ajuba to mitochondria following infection with Sendai virus (Figure 1E), a virus used to trigger the RLR-MAVS pathway. However, we did not observe an interaction between Ajuba and MAVS by immunoprecipitation (IP) (data not shown), suggesting that a role for Ajuba may be indirect. Indeed, ectopic expression of Ajuba in HEK293T cells also reduced the endogenous expression of an additional integral mitochondrial protein, TIMM44. Ajuba-induced loss of both MAVS and TIMM44 was recovered in the presence of either bafilomycin A1 (BafA1) or epoxomicin, inhibitors of lysosome- and proteasome-dependent degradation, respectively (Figure 1D). The major pathway that utilizes both of these cellular degradation pathways is mitophagy. Here, the proteasome degrades outer membrane mitochondrial proteins ubiquitinated by Parkin, whereas lysosomes fused to autophagosomes ultimately degrade damaged mitochondria (Harper et al., 2018; Sekine and Youle, 2018). Consistent with a role in mitophagy, Ajuba expression did not alter MAVS expression in HeLa cells that are naturally deficient in Parkin (Heo et al., 2015; Ordureau et al., 2014). MAVS degradation was restored following expression of wild-type Parkin, but not the C431F active site mutant (Figure 1E). As MAVS signaling is negatively regulated by mitophagy (He et al., 2019; Yang et al., 2018), these results raise the possibility that Ajuba regulates mitophagy more generally rather than having a specific role in RLR-MAVS antiviral responses.

**Figure 1:**
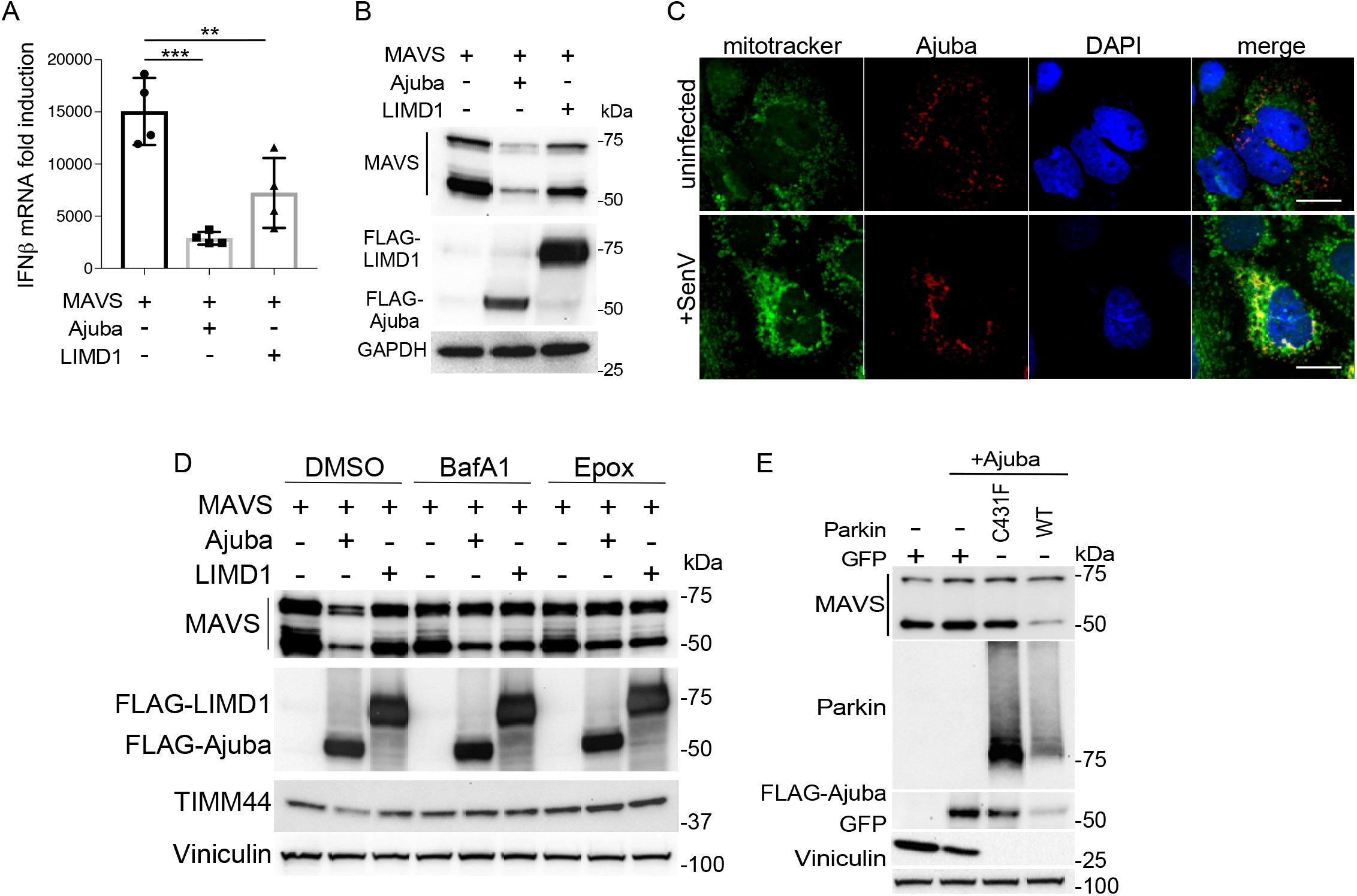
Ajuba negatively regulates MAVS expression. **A.** IFNβ mRNA induction following ectopic expression of MAVS alone or together with Ajuba or LIMD1 in HEK293T cells. **B.** Protein expression from an experiment in A. **C.** Confocal images of FLAG-Ajuba in either mock infected or SenV-infected cells stained for total mitochondria using mitotracker (green) and Ajuba (red). DAPI counterstains the nuclei. **D.** The experiment in A. was repeated with cells treated with 200nM bafilomycin A1 (BafA1) or 200nM epoxomicin (epox) for 6 h prior to cell lysis. **E.** Western blot of HeLa cells transfected with MAVS, WT Parkin, C431F Parkin or Ajuba. DNA was normalized using a GFP expressing control plasmid. Error bars represent mean±SD; *P<0.05, **P<0.01, ***P<0.001 by one-way ANOVA.

### Ajuba is regulated by PINK1-Parkin-mediated degradation during mitophagy

To determine if Ajuba responds generally to mitochondrial damage, mitophagy competent HEK293T cells expressing Ajuba were treated with carbonyl cyanide m-chlorophenyl hydrazone (CCCP) that acts as a proton ionophore to depolarize mitochondrial membrane potential (Fujimaki et al., 2018). CCCP treatment induced translocation of Ajuba from the cytosol to mitochondria-enriched fractions in less than 30 min post treatment (Figure 2A). Similar to overexpression of MAVS, treatment of cells with additional inducers of oxidative stress, tunicamycin and oligomycin, also resulted in reduced expression of integral mitochondrial proteins Mfn1 and TIMM44 in the presence of Ajuba, but not the ER-resident calreticulin (Figure 2B). This suggests that Ajuba responds generally to oxidative stress to specifically promote mitochondrial degradation. In Figures 1E and 2B, we also observed reduced expression of Ajuba when mitophagy was induced for periods longer than 6 h. To examine this more closely, HEK293T cells expressing Ajuba were treated with CCCP or starved to induce bulk autophagy for 6 h and treated with inhibitors of degradation. The CCCP-induced reduction of Ajuba and Mfn1 was dependent on both lysosome- and proteasome-mediated degradation. However, expression of Ajuba or Mfn1 was not responsive to bulk autophagy as evidenced by starvation-induced degradation of the ER-resident autophagy receptor, FAM134B (Khaminets et al., 2015) that is rescued by lysosome inhibition only (Figure 2C). Thus, Ajuba responds specifically to mitochondrial damage. To determine if Ajuba degradation during mitophagy was dependent on PINK1 or Parkin, Ajuba was expressed in the well-established HeLa cell model lacking either protein (Heo et al., 2015). Both basal expression and CCCP-dependent degradation of Ajuba were dependent on PINK1 and Parkin, with Ajuba degradation responding to CCCP and the presence of PINK1 or Parkin in a manner similar to Mfn1 (Figure 2D). To determine if Ajuba has a role in mitochondrial function, we generated an Ajuba^−/−^ mouse model (Supplementary Figure 2), and measured oxygen consumption rates (OCR) by Seahorse Bioanalyzer assays in age-matched WT, Ajuba^−/−^ or PINK1^−/−^ mouse embryonic fibroblasts (MEFs) (Figure 2E, F). Measures of basal respiration ATP-linked respiration, proton leak, spare respiratory capacity and non-mitochondrial respiration were similar between the three MEF genotypes. However, Ajuba^−/−^ MEFs demonstrated increased maximal respiration, thereby confirming that Ajuba regulates mitochondrial function. Taken together, these results suggest that Ajuba responds to a variety of mitochondrial stressors by translocating to mitochondria, where it is subsequently degraded during PINK1/Parkin-dependent mitophagy.

**Figure 2:**
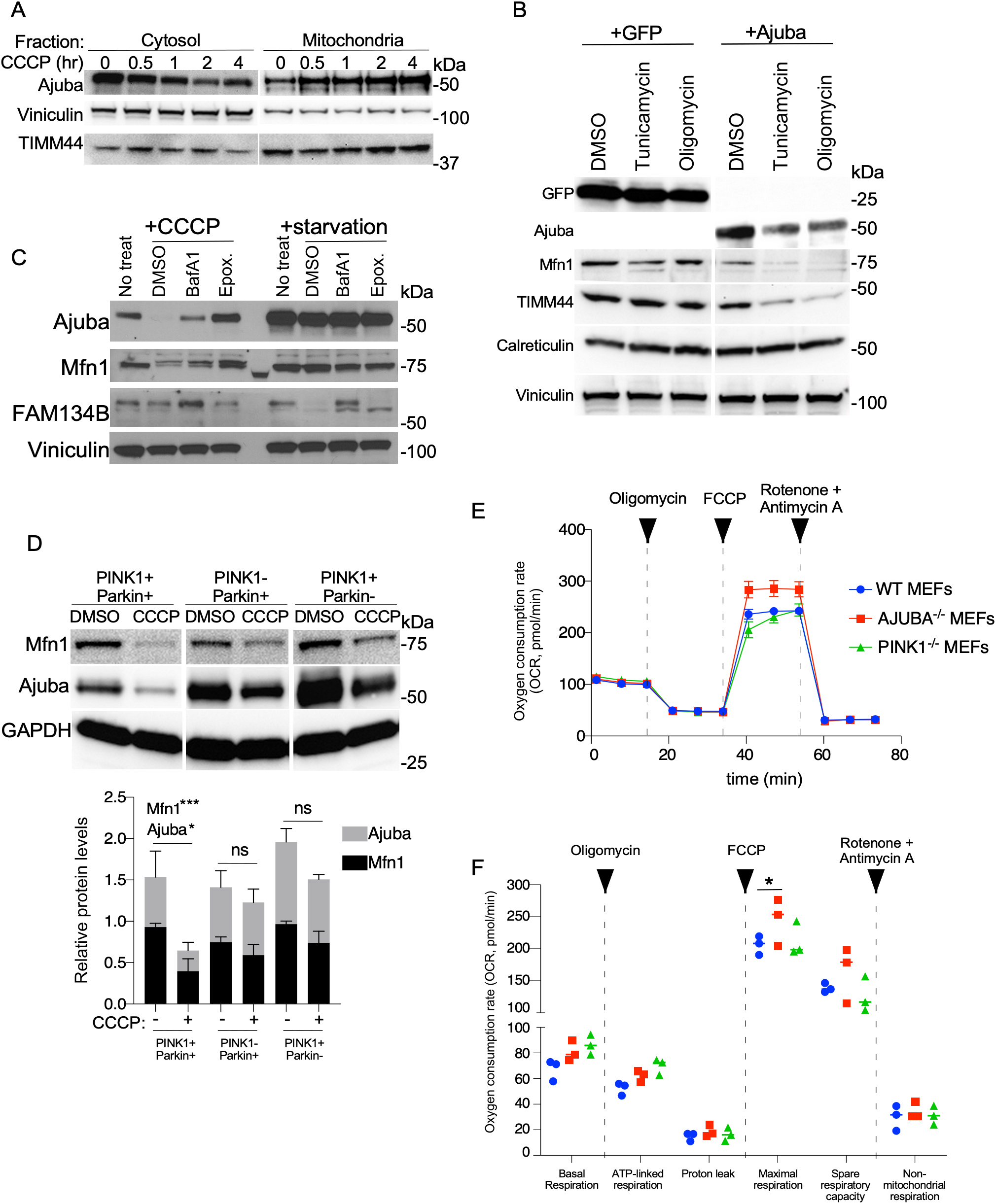
Ajuba is regulated by PINK1-Parkin mediated degradation during mitophagy. **A.** Western blot of HEK293T cells expressing FLAG-Ajuba and treated with 10μM CCCP for the times indicated followed by cellular fractionation. Viniculin and TIM44 are used as fractionation controls for cytosol and mitochondrial enrichment, respectively. **B.** Western blot of HEK293T cells expressing GFP or FLAG-Ajuba and treated with tunicamycin or oligomycin for 5 h at 5μg/ml. **C.** Western blot of HEK293T cells expressing FLAG-Ajuba following treatment with CCCP or starvation in Earl’s balanced salt solution (EBSS) for 6 h. Mfn1 and FAM134B are used as positive controls for mitophagy and bulk autophagy, respectively. **D.** HeLa cells engineered to express Parkin or knocked out for PINK1 were transfected with FLAG-Ajuba and treated with 10μM CCCP for 6 h. Quantification of Ajuba and Mfn1 expression by densitometry for 3 experiments is shown. **E.** Representative Seahorse analysis of oxygen consumption rate (OCR) in WT, Ajuba^−/−^ or PINK1^−/−^ MEFs following treatment with oligomycin (2μM), FCCP (2μM) or rotenone + Antimycin A (0.5μM each). **F.** Calculated OCR associated with mitochondrial functions from three independent experiments performed in triplicate (mean±SD); *P<0.05 by two-way ANOVA.

### Ajuba interacts with PINK1 to promote PINK1 autophosphorylation and mitophagy

To determine if Ajuba is required for mitophagy, Ajuba ^−/−^ MEFs were labeled simultaneously with Mitotracker red and Mitotracker green, treated with CCCP, and the quenching of green fluorescence following acidification in lysosomes was measured by flow cytometry (Sprung et al., 2018). Mitochondrial mass was not different in resting MEFs as measured by Mitotracker green mean fluorescence intensity (MFI) (Figure 3A). However, as indicated by the shift in ratio of red:green MFI following CCCP treatment, mitophagy was reduced in Ajuba^−/−^ MEFs, and not different from WT MEFs treated with CCCP and BafA1 to inhibit lysosomal acidification (Figure 3B). This finding suggests that depolarized mitochondria are not delivered efficiently to lysosomes in the absence of Ajuba.

**Figure 3:**
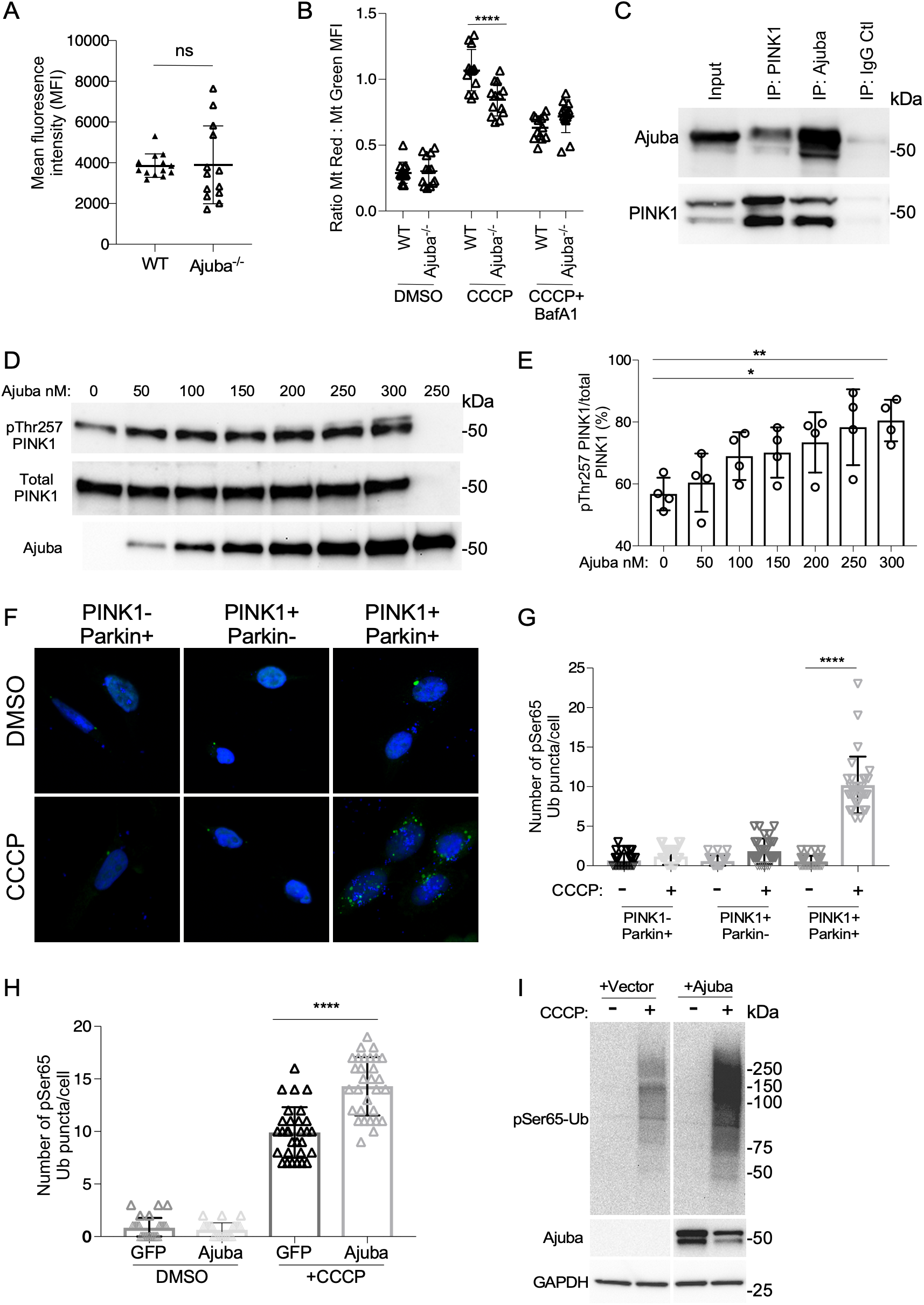
Ajuba interacts with PINK1, and promotes PINK1 autophosphorylation and mitophagy. **A.** Mean fluorescence intensity of WT or Ajuba^−/−^ MEFs were labeled with mitotracker green (200nM). **B.** WT or Ajuba^−/−^ MEFs were labeled with mitotracker green and mitotracker red (200nM), treated with 10μM CCCP for 16 h with or without BafA1, and analyzed by flow cytometry. **C.** Reciprocal IP of *E.coli*-expressed human PINK1 and Ajuba. **D.** Autophosphorylation of human PINK1 (300nM) in the presence of increasing concentrations of Ajuba (0-300nM). Total protein was normalized with bovine serum albumin. **E.** Quantification of pThr257/total PINK1. **F.** Confocal microscopy of HeLa cells transfected with a plasmid expressing HA-Ub, followed by treatment with 10μM CCCP for 2 h and stained for pSer65-Ub (green). Nuclei were counterstained with DAPI (blue). **G.** Quantification of pSer65-Ub puncta per cell in HeLa cells in E demonstrating that signals are specific to PINK1-Parkin mediated mitophagy. **H.** pSer65-Ub assay in HEK293T cells expressing GFP or Ajuba. **I.** Western blot for total pSer65-Ub in HEK293T cells expressing Ajuba and treated with 10μM CCCP for 2 h. Error bars represent mean±SD from 3-4 independent experiments; *P<0.05, **P<0.01, ***P<0.001 by one-way ANOVA.

**Figure 4:**
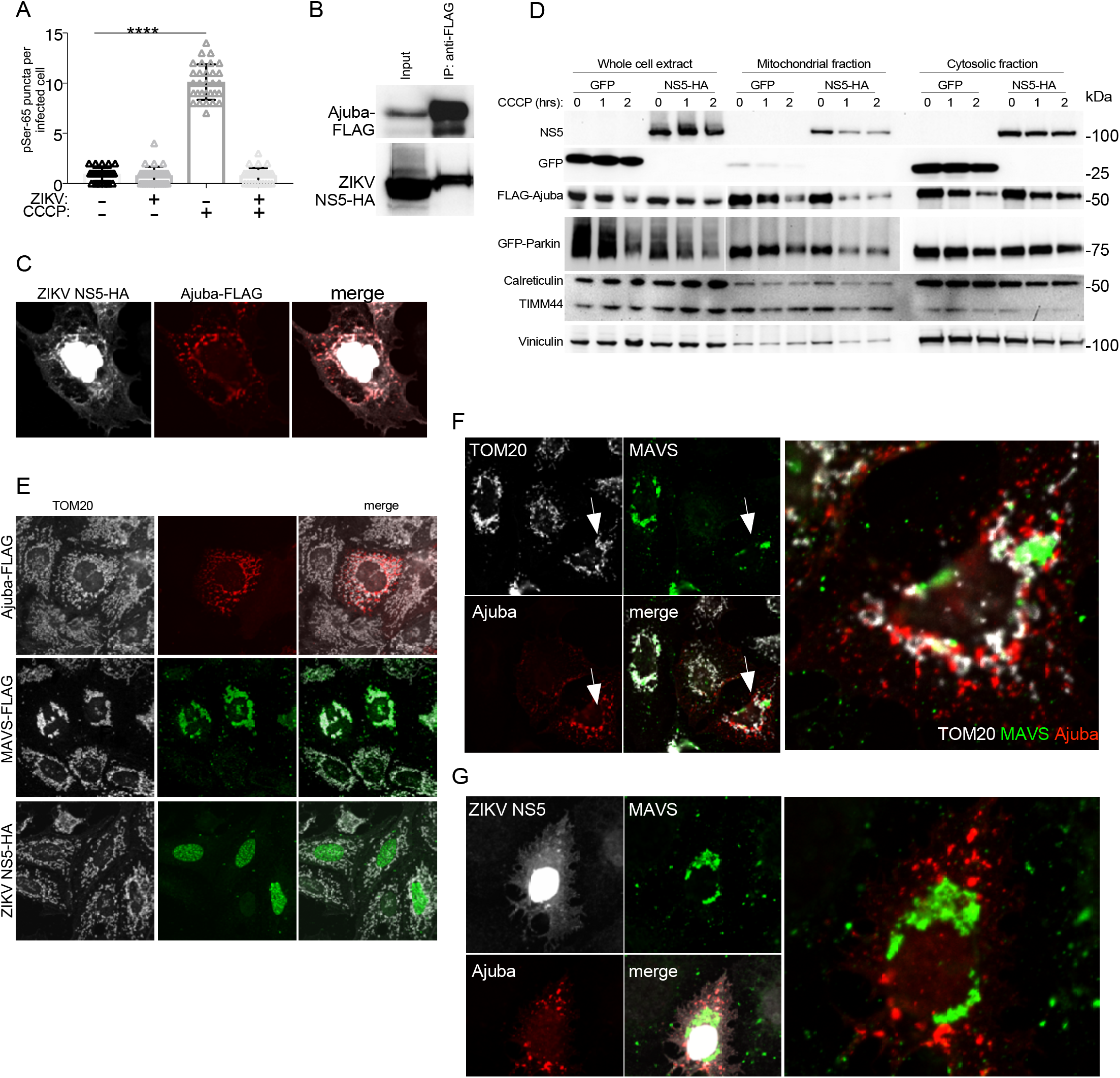
Zika virus NS5 binds to Ajuba to suppress mitophagy. **A.** pSer65-Ub assay in HEK293T cells infected with ZIKV for 48 h and then treated with or without 10μM CCCP for 2 h. Error bars represent mean±SD; ***P<0.001 by one-way ANOVA. **B.** IP of FLAG-Ajuba and ZIKV NS5-HA expressed in HEK293T cells. **C.** Confocal microscopy demonstrating cytosolic co-localization of FLAG-Ajuba (red) and ZIKV NS5-HA (greyscale). **D.** HEK293T cells were transfected with plasmids expressing FLAG-Ajuba, GFP-Parkin and either GFP or ZIKV NS5-HA. Cells were treated with 10μM CCCP for the times indicated. Cells were fractionated into mitochondrial fractions (enriched for TIMM44) or cytosolic fractions (enriched for viniculin and calreticulin) demonstrating that CCCP-induced mitochondrial localization of Ajuba and Parkin is delayed in the presence of ZIKV-NS5. **E.** Confocal microscopy of Huh7 cells expressing single plasmids encoding Ajuba-FLAG, MAVS, or ZIKV NS5-HA. Only MAVS strongly colocalizes with mitochondria stained for TOM20 (greyscale). **F.** Confocal image of cells co-expressing MAVS and Ajuba demonstrating recruitment of Ajuba to MAVS-positive mitochondria and disruption of MAVS aggregates (arrow). **G.** Confocal image demonstrating that co-expression of ZIKV NS5 in cells also expressing MAVS and Ajuba suppresses Ajuba recruitment and maintains MAVS aggregates.

To determine a functional role for Ajuba in PINK1-Parkin signaling, we turned to the characterized cellular role of Ajuba in relief of kinase autoinhibition, particularly mitotic kinases such as Aurora-A (Kashatus et al., 2011). Similar to these mitotic kinases, human PINK1 is a Ser/Thr kinase whose resting state is in an autoinhibited conformation. However, the mechanism(s) by which this autoinhibition is relieved following stabilization on the outer mitochondrial membrane is not completely characterized (Harper et al., 2018). We hypothesized that once at the mitochondria, Ajuba may bind to PINK1 and function in its recognized role of kinase activation. Ajuba and PINK1 expressed from *E. coli* interacted directly with each other by reciprocal co-IP (Figure 3C), a finding that was supported by co-localization of the two proteins in cells by immunofluorescence assay (IFA) (Supplemental Figure 3A). Importantly, *E.coli*-expressed PINK1 has low autophosphorylation activity (Rasool et al., 2018), but auto-phosphorylation of PINK1 at Thr257 significantly increased when PINK1 was incubated with increasing levels of Ajuba (Figure 3D,E). Ajuba can be phosphorylated by the kinase with which it interacts (Chen et al., 2016) prompting us to test if PINK1 phosphorylates Ajuba following coexpression by IP and mass-spectrometry. Based on the PINK1 phosphorylation consensus sequence (Torii et al., 2020), two residues in the pre-LIM domain of Ajuba are predicted to be phosphorylated by PINK1, S39 and S136. Coexpression of PINK1 with Ajuba resulted in a higher molecular mass form of Ajuba (Band ‘B’; Supplemental Figure 3B). Peptides visible by mass spectrometry spanned an average of 48% of Ajuba (range 29-63%) and included both of these residues, but only S39 was specifically phosphorylated in the presence of PINK1 and not TBK1, another Ser/Thr kinase used as a control (Supplemental Figure 3B). These results demonstrate direct and functional interactions between Ajuba and PINK1.

To determine if Ajuba affects PINK1 function in cells, we developed an IFA to quantify pSer65-Ub in individual cells. HeLa cells were transfected with HA-Ub, treated with CCCP and stained for pSer65-Ub. The number of puncta were then counted per cell. The assay was validated in HeLa cells lacking either PINK1 or Parkin, demonstrating that pSer65-Ub puncta were dependent on both proteins and coupled to mitochondrial depolarization (Figure 3F,G). Ajuba expression did not induce spontaneous pSer65-Ub puncta, but did increase pSer65-Ub accumulation in CCCP-treated cells compared to a GFP control (Figure 3H). In addition, ectopic expression of Ajuba increased the total level of pSer65-Ub in CCCP-treated cells (Figure 3I). Taken together, these results suggest that Ajuba is required for efficient mitophagy and functions by directly interacting with PINK1 to promote pSer65-Ub accumulation.

### Zika virus NS5 binds to Ajuba to suppress mitophagy

To examine the role of Ajuba in mitophagy further, we first determined the effect of ZIKV replication on mitophagy. CCCP-induced pSer65-Ub puncta were markedly reduced in ZIKV-infected cells, suggesting that ZIKV exerts a strong block to mitophagy (Figure 4A). To determine if this was associated with NS5 binding to Ajuba, we first confirmed the interaction. IP of Ajuba resulted in coprecipitation of ZIKV NS5 in HEK293T cells (Figure 4B). ZIKV NS5 localizes mainly to the nucleus, but low levels of cytosolic NS5 could be observed co-localizing with Ajuba following co-expression (Figure 4C). Consistent with viral inhibition of mitophagy, ZIKV NS5 reduced accumulation of Ajuba as well as Parkin (used as a downstream target of PINK1) in the mitochondrial fraction following treatment of cells with CCCP (Figure 4D). We also examined localization of Ajuba by IFA with and without NS5 when using MAVS as a stimulus for mitochondrial depolarization. When expressed alone, MAVS localized to mitochondria and caused mitochondrial aggregation (Figure 4E). When Ajuba and MAVS were coexpressed in the same cell, Ajuba appeared to interweave with the mitochondrial marker TOM20 and disrupt aggregates of MAVS (Figure 4F). However, coexpression of ZIKV NS5 suppressed mitochondrial recruitment of Ajuba in MAVS-positive cells (Figure 4G). These results suggest that ZIKV imparts a remarkable block to mitophagy that is mediated by NS5 binding to Ajuba to suppress its recruitment to depolarized mitochondria.

#### Suppression of mitophagy results in mtRNA release, PKR phosphorylation and PKR-dependent chemokine expression

Perhaps one of the most significant consequences of viral antagonism of mitophagy and failure to clear damaged mitochondria from infected cells is the potential to amplify inflammatory responses (Moehlman and Youle, 2020). To determine the consequence of the viral block in mitophagy, we first performed RNAseq in ZIKV-infected WT, Ajuba^−/−^ or PINK1^−/−^ MEFs. In both mock- and ZIKV-infected cells, the majority of differentially regulated genes (DEGs) were commonly observed in PINK1^−/−^ and Ajuba^−/−^ cells, providing additional evidence that the two proteins function in the same pathway (Figure 5A,B). Notably, ZIKV-infected PINK1^−/−^ and Ajuba MEFs^−/−^ demonstrated increased activation of antiviral and inflammatory pathways (Supplementary Figure 4A). Specifically, chemokines recognized as hallmarks of human infection with ZIKV (Foo et al., 2018; Kam et al., 2017; Michlmayr et al., 2020) were increased by 72 hpi in the absence of Ajuba or PINK1 (Figure 5C and Supplemental Figure 4B). Increased chemokine expression appeared independent of substantial increases in virus replication even when IFN signaling was blocked, as measured by release of infectious virus (Figure 5D,E).

**Figure 5:**
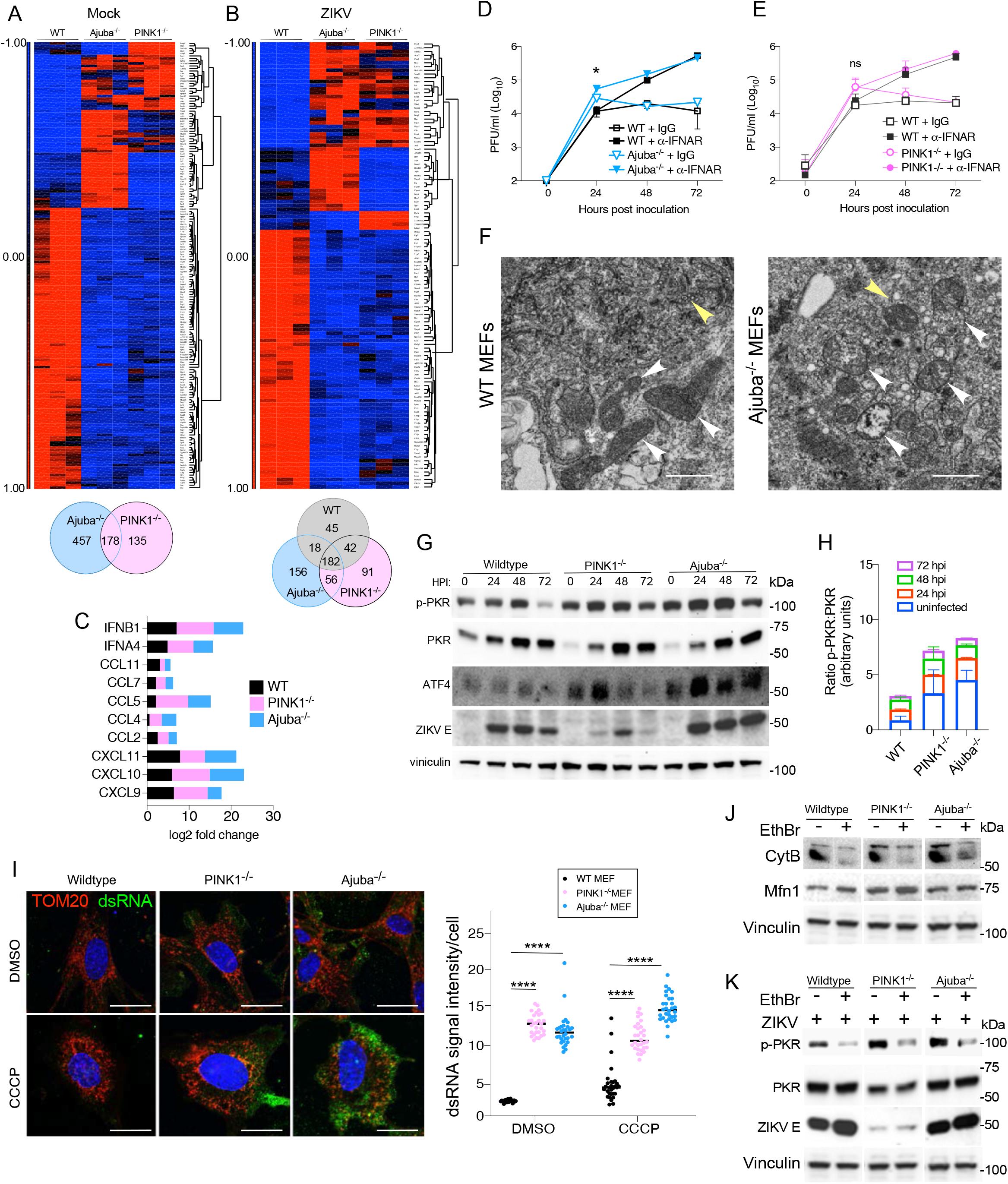
ZIKV replication in mitophagy-deficient MEFs is associated with amplified chemokine expression, PKR phosphorylation, and release of mitochondrial dsRNA. **A.-B.** RNAseq in 3 replicate cultures of WT, PINK1^−/−^ or Ajuba^−/−^ MEFs that were **A.** mock-infected, or **B.** infected with ZIKV with MOI 0.1 and harvested at 72 hpi. **C.** Chemokine and IFN gene expression changes (log2 fold change) in ZIKV-infected cells compared to mock-infected cells at 72 hpi. **D.** ZIKV growth curves in WT (black) and Ajuba^−/−^ (blue) MEFs treated with isotype control (open symbols) or anti-IFNAR1 mAb (closed symbols). *P≦0.05 by Mann-Whitney test. **E.** ZIKV growth curves in WT (black) and PINK1^−/−^ (pink) MEFs treated with isotype control (open symbols) or anti-IFNAR1 mAb (closed symbols). **F.** Transmission electron microscopy images of WT and Ajuba^−/−^ MEF at 72 hpi with ZIKV. White arrowheads indicate mitochondria; yellow arrowheads indicate sites of virus replication. **G.** Western blot demonstrating that ZIKV infection increases expression levels of PKR, phospho-PKR and ATF4 in the absence of PINK1 or Ajuba. Basal levels of phospho-PKR and ATF4 are also increased in uninfected PINK1^−/−^ and Ajuba^−/−^ MEFs compared to WT MEFs. **H.** Quantification by densitometry of p-PKR/PKR ratio in ZIKV-infected MEFs from 2 independent experiments. **I.** WT, PINK1^−/−^ or Ajuba^−/−^ MEFs were stained for mitochondria (TOM20, red) and dsRNA (J2, green) at 4 h post treatment with vehicle or CCCP. Staining intensity of dsRNA was quantified by image J software . ****P<0.001 by two-way ANOVA. **J.** Treatment of MEFs with ethidium bromide (EthBr) demonstrating loss of mitochondrial genome-encoded cytochrome B (cytochrome B) but not nuclear genome-encoded Mfn1. **K.** EthBr treatment suppresses PKR phosphorylation in ZIKV-infected MEFs.

The lack of major differences in viral replication suggests that increased chemokine responses may result from mitochondrial DAMPs. Compared to WT MEFs, mitochondria in ZIKV-infected Ajuba^−/−^ MEFs contained disorganized cristae and loss of double membrane structure suggesting considerable damage (Figure 5F). We also noted from the RNAseq data upregulation of the pathway ‘Role of PKR in IFN induction and antiviral response’ in ZIKV-infected Ajuba- and PINK1-knockout MEFs (Supplementary Figure 4B, C). PKR is a stress-associated kinase that binds to double stranded RNA (dsRNA) of viral or mitochondrial origin (Kim et al., 2018), or it can be separately activated as part of the integrated stress response (ISR) (Hou et al., 2017). Activated PKR then phosphorylates eIF2α to induce expression of the stress associated transcription factor, ATF4 (Pakos-Zebrucka et al., 2016). In support of the RNAseq data, PKR phosphorylation and ATF4 expression were elevated basally in uninfected Ajuba^−/−^ and PINK1^−/−^ MEFs, and further increased by ZIKV infection (Figure 5 G,H). Mitochondrial RNA (mtRNA) can be visualized using the J2 mAb with high specificity (95-99%) in cultured cells (Dhir et al., 2018). Compared to WT MEFs, mtRNA staining was both increased in intensity and localized outside of mitochondria in the absence of either Ajuba or PINK1 even without additional stimulus for depolarization by CCCP treatment (Figure 5I). To determine if PKR is activated by mtRNA, cells were treated with ethidium bromide (EthBr) to deplete mitochondrial RNA as evidenced by reduced expression of the mitochondrially encoded cytochrome B, but not the nuclear encoded Mfn1. Depletion of mtRNA also suppressed PKR phosphorylation without affecting total PKR expression in ZIKV-infected MEFs (Figure 5 J,K). Taken together, these results provide evidence for release of mtRNA to the cytosol when mitophagy is compromised in primary cells resulting in PKR activation.

To understand how ZIKV replication, PKR activation, and cytokine expression are linked, we infected primary human dermal fibroblasts with ZIKV and treated the cells with inhibitors of PKR (C16) or eIF2α (ISRIB) phosphorylation (Gal-Ben-Ari et al., 2018; Rabouw et al., 2019). Both inhibitors reduced virus replication consistent with the established roles of PKR in phosphorylating eIF2α (Gal-Ben-Ari et al., 2018) and the known pro-viral role of stress responses that drive eIF2α phosphorylation (Ambrose and Mackenzie, 2011, 2013; Hou et al., 2017)(Figure 6A-C). However, we were surprised to observe that expression of multiple chemokines including CXCL10, CXCL1, CXCL12, CCL2, CCL4, and CCL5, as well as select cytokines (IL-1α and IL-18) were highly dependent on PKR but not eIF2α (Figure 6D-E). Indeed, inhibition of eIF2α-phosphorylation exacerbated chemokine and cytokine expression, consistent with failure to resolve cellular stress through the ISR. Importantly, as virus replication was reduced in both C16- and ISRIB-treated cells, these findings uncouple virus replication from the inflammatory response and support the finding that mtRNA/DAMP signaling is a significant driver of PKR-dependent inflammation. Our data suggests that ZIKV suppression of mitophagy amplifies the ISR which is favorable for flavivirus replication. However, PKR activation in response to mitochondrial damage serves to amplify the pro-inflammatory chemokine response that is the hallmark of flavivirus infection.

**Figure 6:**
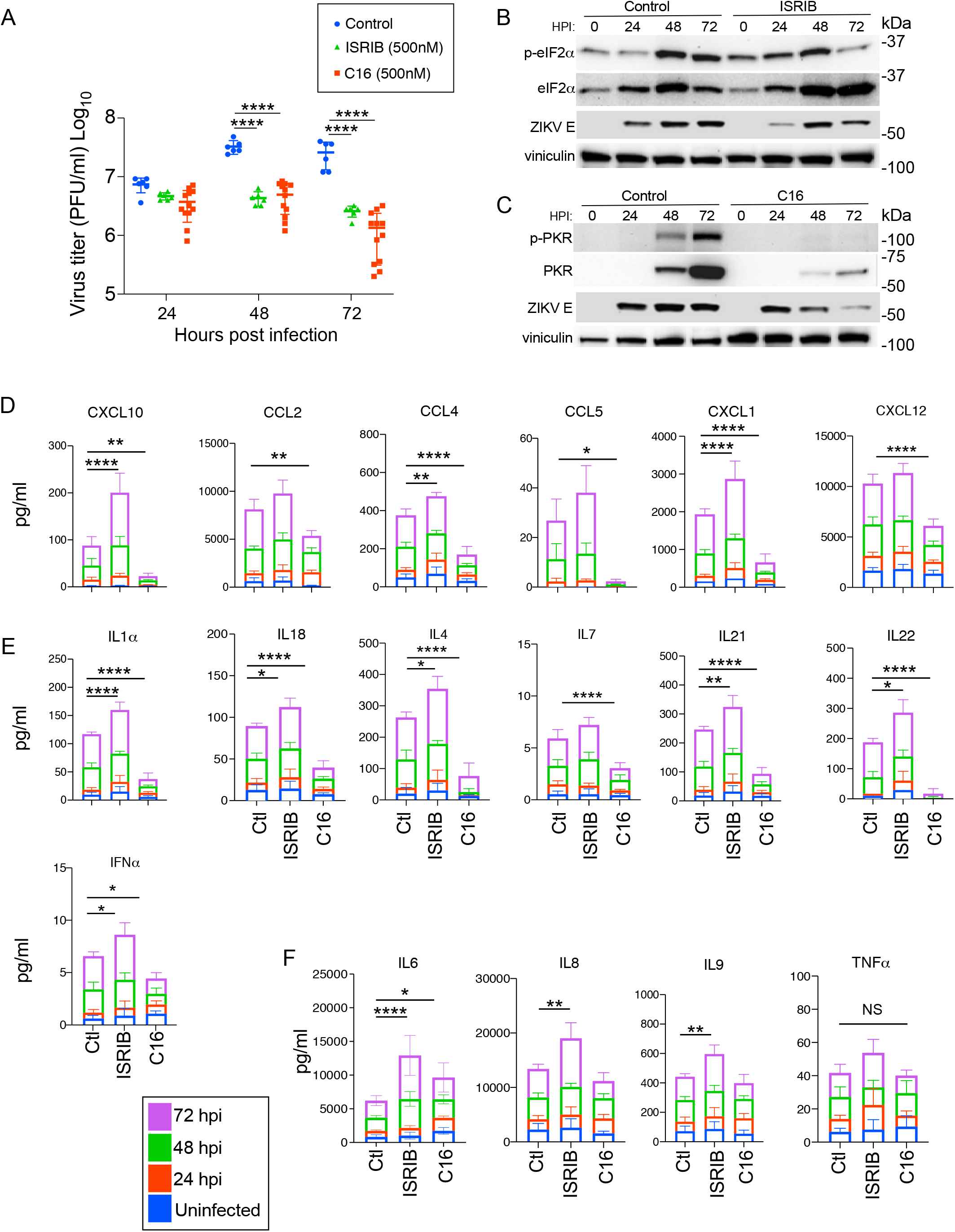
The ISR favors virus replication but is uncoupled from pro-inflammatory chemokine and cytokine expression dependent on PKR. Primary human dermal fibroblasts were infected with ZIKV (MOI 1) and treated with 500nM ISRIB or C16 for 24h prior to harvest. Release of infectious virus was measured by plaque assay. Inhibition of **B.** eIF2α phosphorylation or **C.** PKR phosphorylation was verified by Western blotting. **D.** Chemokines and **E-F.** cytokines were measured by Bioplex multiplex assay in culture supernatants from uninfected cells or cells infected with ZIKV at 24, 48 and 72 hpi. Error bars represent mean±SD from two experiments performed in triplicate; *P<0.05, **P<0.01, ***P<0.001, ****P<0.0001 by one-way ANOVA.

#### Pro-inflammatory chemokines are expressed earlier in ZIKV-infected Ajuba^−/−^ mice associated with increased virus dissemination to tissues

To determine the consequences of suppressed mitophagy in vivo, we examined the kinetics of induction of pro-inflammatory cytokines in Ajuba-deficient mice. Using RNAscope assays, *Ajuba* mRNA expression was particularly enriched in epithelial cells of the skin (keratinocytes) and testes (Sertoli cells), and in epithelial and endothelial cells of the lung. *Ajuba* was also expressed in hepatocytes and endothelial cells of the liver, and ependymal cells, neurons, and epithelial cells in the CNS (Supplementary Figure 5). Thus, *Ajuba* is expressed in flavivirus target cells and organs. To determine if effects on inflammation can be observed in vivo, WT and Ajuba^−/−^ mice were treated with anti-IFNAR (MAR1) mAb one day prior to footpad inoculation with ZIKV to examine responses in the absence of IFN as a confounding factor in mouse models of ZIKV infection (Gorman et al., 2018; Grant et al., 2016). Viremia was equivalent at 3 and 5 dpi (Figure 7B). However, acute response chemokines were either significantly elevated (CXCL1, CXCL10, CCL7) or trending higher (CCL5, CXCL2) in serum at 3dpi (Figure 7A). In contrast, production of late response cytokines like IFNγ were not affected (Figure 7A). We next examined virus burden in tissues. This revealed the striking finding that virus titers in target tissues including the spleen and brain were up to 80-fold higher at 3dpi in Ajuba^−/−^ mice, despite no differences in viremia. However, by 5 dpi, ZIKV titers in tissues were generally not different between WT and Ajuba^−/−^ mice, demonstrating that there is no intrinsic advantage to virus replication (Figure 7B). These results suggest that suppression of mitophagy by ZIKV facilitates virus invasion of tissue. The pre-treatment of mice with anti-IFNAR mAb suggests that the difference in cytokine expression and virus dissemination at 3 dpi is largely independent of type I or III IFNs, and the equivalent viremia suggests that elevated chemokine expression in serum is not driven by increased viral pattern-associated molecular patterns (PAMPs). Instead, these findings suggest that inhibition of Ajuba and suppressed mitophagy by ZIKV amplifies early expression of acute response chemokines in vivo and that this inflammation is strongly linked to DAMP signaling and virus invasion of tissues.

**Figure 7:**
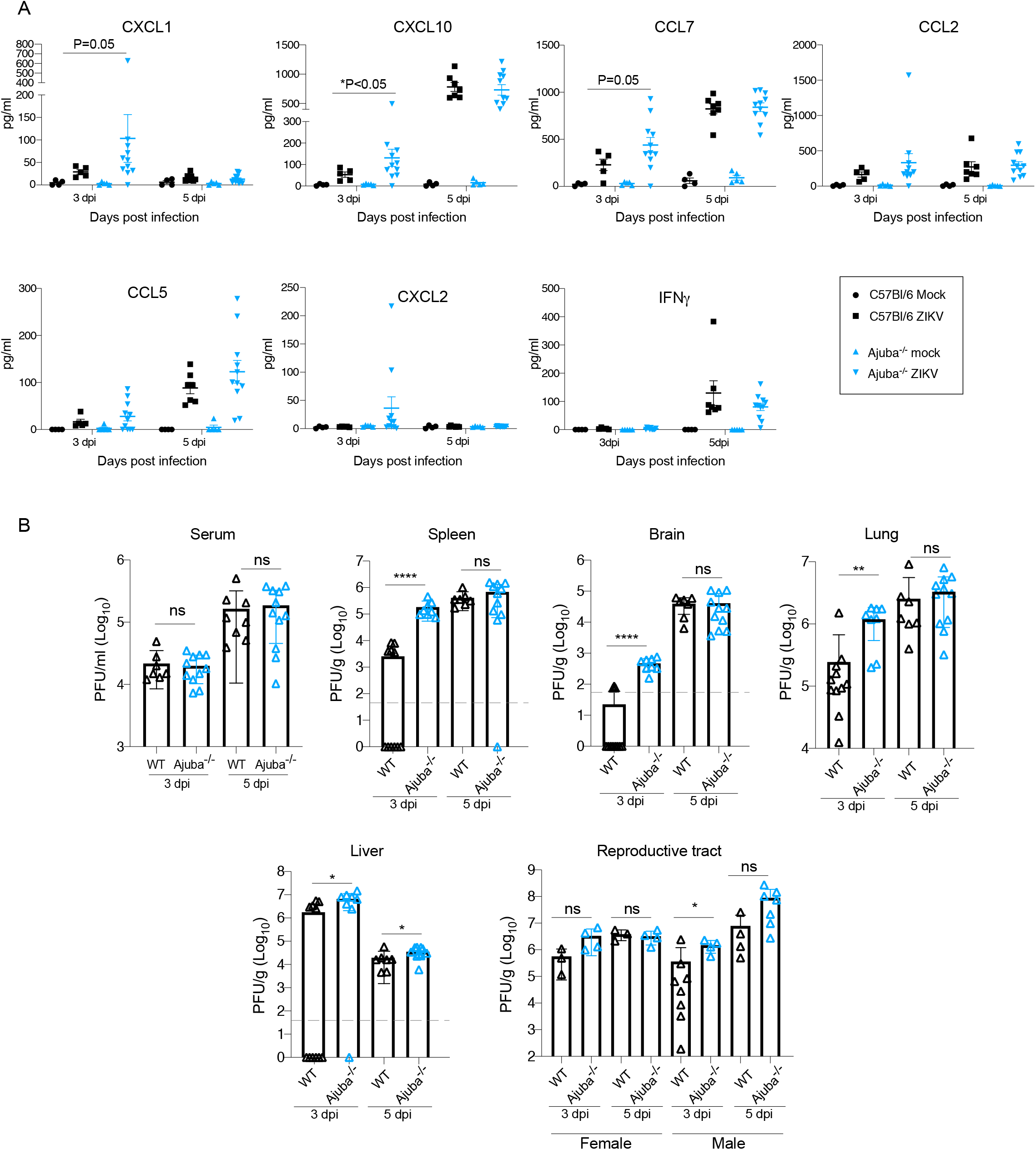
Pro-inflammatory chemokines are expressed earlier in ZIKV-infected Ajuba^−/−^ mice associated with increased virus invasion of tissues. WT and Ajuba^−/−^ mice were treated with 2 mg anti-IFNAR mAb one day prior to infection with 10^3^ PFU ZIKV. **A.** Serum protein levels of the chemokines and cytokines measured by Bioplex multiplex assays. *P≦0.05 by one-way ANOVA. **B.** Virus titers at 3 and 5 dpi in serum and tissues measured by plaque forming units (PFU)/ml serum. Error bars represent mean±SD; *P<0.05, **P<0.01, ****P<0.0001 by Mann-Whitney test.

## DISCUSSION

By studying virus-host interactions, this work identifies Ajuba as a critical regulator of PINK1-Parkin-dependent mitophagy. Ajuba participated in mitophagy following multiple stressors, including ER-stress, MAVS activation, or direct mitochondrial depolarization, suggesting that Ajuba is generally important to mitochondrial quality control. ZIKV suppressed PINK1-Parkin-dependent mitophagy early in the pathway prior to the accumulation of pSer65-Ub, consistent with a role for Ajuba in positive regulation of PINK1 activation. PINK1 activation following accumulation on mitochondrial membranes results from self-phosphorylation of the activation loop *in trans* presumably as the local concentration of PINK1 increases (Sekine and Youle, 2018). However, we provide several lines of evidence that Ajuba may function to further augment PINK1 activation including demonstration of a direct interaction between PINK1 and Ajuba, augmentation of both PINK1 autophosphorylation in vitro and pSer65-Ub in cells by Ajuba, and RNAseq data suggesting that PINK1 and Ajuba function in overlapping regulatory cascades. While the most widely conserved regulatory mechanism for Ser/Thr kinase activation is phosphorylation of kinase activation loop residues, stabilization of conformational changes occurs following kinase binding to cofactors that independently increase kinase activity by up to several hundred-fold (Dodson and Bayliss, 2012; Zorba et al., 2014). Additional careful biochemical studies are needed to determine the precise relationship between PINK1 and Ajuba. However, the identified interaction suggests a potential mechanism for further stabilization of PINK1 conformation to ensure a regulated acceleration of kinase activity, as observed for Aurora-A (Ruff et al., 2018). Further definition of Ajuba’s role in PINK1-Parkin signaling may reveal new avenues for therapeutic interventions in PD.

The consequences of damaged mitochondria to inflammation is currently of intense scientific interest due to the central role of these processes in neurological disorders (e.g. PD, and AD) and cellular transformation. The mechanisms of inflammation are linked to the ancient origins of mitochondria as α-proteobacterium, and are mediated through DAMP activation of innate immune signaling pathways (Mottis et al., 2019; Youle, 2019). For example, mitochondrial DNA is sensed through cGAS-STING, a process that is amplified in PINK1^−/−^ or PARK2^−/−^ deficient mice following metabolic stress (Sliter et al., 2018). mtRNA is also released and can be sensed by PKR or Mda5 (Dhir et al., 2018; Kim et al., 2018). PKR activation by mtRNA and downstream eIF2α phosphorylation has been shown, but role for PKR sensing of mitochondrial health as a central amplifier of chemokine expression during virus infection has not been previously reported. Here, by investigating interactions between ZIKV and mitochondria, we reveal that viral antagonism of mitophagy is directly translated to inflammation through PKR-sensing of mtRNA. Moreover, we show that viral antagonism of mitophagy is a key determinant of virus invasion of tissues linked to pro-inflammatory chemokine responses.

Pro-inflammatory chemokine expression is linked to disease severity in humans infected with ZIKV, DENV and YFV (Foo et al., 2018; Kam et al., 2017; Michlmayr et al., 2017; Michlmayr et al., 2020; Naveca et al., 2018) although the molecular mechanisms leading to their initial expression are not well defined. As chemokines are ultimately responsible for coordinating leucocyte recruitment and activation, determining the early events that drive chemokine expression is a key question in flavivirus pathogenesis. Here we reveal that pro-inflammatory chemokine responses are amplified in ZIKV-infected cells through PKR sensing of damaged mitochondria that are actively retained in cells through viral inhibition of mitophagy. CXCL10 and CCL2 have been specifically linked to symptomatic ZIKV infection during pregnancy and congenital abnormalities (Foo et al., 2018; Kam et al., 2017; Naveca et al., 2018). In addition, CXCL10, CCL2, CCL5 and IL-10 have been identified as the central cytokines connecting temporal changes in cell populations and gene expression in the context of ZIKV-infected Nicaraguan children (Michlmayr et al., 2017; Michlmayr et al., 2020). Strikingly, compared to WT mice, Ajuba^−/−^ mice exhibited an earlier amplification of these same cytokines including CXCL10, CCL2, CCL5, and CXCL1 that were similarly elevated at early-acute times in the Nicaraguan cohort, with CCL2, and CXCL10 remaining high during late-acute phase of infection in humans and in our mouse model. Severe disease following flavivirus infection generally occurs after peak virus burden in blood (Tricou et al., 2011). The reasons for this are multifaceted, but our results suggest a specific mechanism for sustained inflammation in tissues due to compromised mitophagy and increased DAMP signaling. As illustrated by infection of primary human fibroblasts, once this signaling is initiated, even a reduction in virus titer through inhibition of the ISR was not sufficient to suppress chemokine expression. Evidence for these processes in humans has been observed in placental tissues from ZIKV-infected mothers including mitochondrial dysfunction, metabolic alterations and induction of inflammatory mediators (Chen et al., 2020). The current work identifies a molecular mechanism for these observations encoded through ZIKV NS5 to amplify mitochondrial stress responses in infected tissue (Supplementary Figure 6). Thus, pathways of mitophagy and DAMP signaling are identified as potential therapeutic targets in severe flavivirus disease.

Multiple viruses increase mitophagy as a strategy to evade the IFN response (Zhang et al., 2018). Thus, the finding that ZIKV inhibits mitophagy to maintain damaged mitochondria in cells and amplify inflammatory signaling cascades is surprising. The observation that ZIKV encodes at least two mechanisms to manipulate mitochondrial dynamics though NS5 (this paper) and NS4B suppression of mitochondrial fission (Chatel-Chaix et al., 2016) suggests that mitochondrial functions are important to virus biology. The pro-viral function of impaired mitophagy will be associated in part with amplification of the ISR as a known factor that favors flavivirus replication through suppression of host mRNA translation (McEwen et al., 2005). However, we observed no fundamental advantage to virus replication in primary MEFs or in mouse tissues lacking Ajuba. Instead, we found that the major consequence of suppressed mitophagy in vivo is to facilitate virus invasion of tissues. The precise events required for flavivirus invasion of critical tissues are not understood (Ayala-Nunez and Gaudin, 2020), although experimental evidence suggests that inflammatory responses could increase virus transmigration across endothelial barriers (Miner and Diamond, 2016). We observed an 80- and 16-fold increase in virus burden in the spleen and brain, respectively, at 3 dpi in Ajuba^−/−^ mice. This increase was independent of viremia levels, intrinsic differences in virus replication, or IFNAR1 signaling, but was associated with earlier expression of chemokines that are considered hallmarks of monocyte recruitment and activation. Monocytes are primary targets for flavivirus infection in the blood (Michlmayr et al., 2017), and ZIKV has been shown to increase monocyte adhesion and transmigration across endothelial barriers (Ayala-Nunez et al., 2019). We therefore speculate that monocyte activation increases tissue seeding of virus. Once in tissues, it is possible that retention of damaged mitochondria in infected cells further promotes tissue injury. To further address these fundamental questions in flavivirus pathogenesis, it will be important to determine the spatial and temporal kinetics of chemokine expression, ZIKV infection and monocyte activation status in blood and key tissues.

In summary, we have shown that ZIKV NS5 antagonizes mitophagy by binding to the host protein Ajuba and thereby prevents Ajuba translocation to depolarized mitochondria where it is required for PINK1-Parkin signaling. The consequences of this to ZIKV include increased DAMP signaling and increased viral dissemination to tissues. It is well established that viral PAMPs initiate pro-inflammatory responses (Gilfoy and Mason, 2007; Samuel et al., 2006). However, our results suggest that mitochondrial stress leading to release of DAMPs is responsible for amplification of specific chemokines following flavivirus infection, and that this is tightly associated with viral invasion of tissues. Suppression of mitophagy also provides a potential selection pressure for development of flavivirus strategies to antagonize IFN responses downstream of mitochondria including suppression of cGAS-STING signaling to mitigate mtDNA release (Aguirre et al., 2017; Aguirre et al., 2012; Yu et al., 2012), and the property of NS5 as a very potent JAK-STAT antagonist to suppress downstream IFN signaling in infected cells (Best, 2017). In addition, ZIKV encodes blocks to RLR-MAVS signaling downstream of MAVS, including suppression of TBK1 and IRF3 activation (Xia et al., 2018). Moreover, these findings raise new potential mechanisms of immune activation and dysfunction in the context of flavivirus infection based on recognized roles of mitophagy in suppressing DAMP-driven responses and autoimmunity (reviewed in (Mottis et al., 2019; Youle, 2019). Finally, our findings illustrate a role of mitophagy in protection from inflammatory responses and virus infection in vivo, beyond the context of neurodegeneration. It is noteworthy that therapeutics aimed at augmentation of mitophagy improve cellular function in animal models of AD (Fang et al., 2019). Thus, therapeutics under development for PD, AD and other disorders of mitochondrial dysfunction should be considered for testing in flavivirus infection models. In this case, augmented mitophagy may dampen pro-inflammatory responses and limit virus seeding of tissues.

## Acknowledgements

This work was supported by the Division of Intramural Research, National Institute of Allergy and Infectious Diseases, National Institutes of Health. Thank you to Stacy Ricklefs and Kimmo Virtaneva (RML RTB, NIAID) for help with RNA extractions, library preparation and sequencing. Thank you to Dr. Wade Harper (Harvard Medical School) for providing the PINK1/Parkin HeLa cell system, Dr. Chengyu Liu (NHLBI Transgenic Core, NIH) for generation of Ajuba^−/−^ mice, and Ryan Kissinger (Visual Medical Arts, RML, NIAID) for graphic arts expertise.

## Author contributions

Conceptualization, S.S.P. and S.M.B; Methodology, S.S.P., S.J.R., K.L.M., F.J., C.S., C.M. and S.M.B.; Investigation, S.S.P., S.J.R., K.L.M., G.L.S., M.L., F.J., C.K., D.G., A.H., C.S., R.R., G.S., and C.M.; Resources, C.M.B, G.S., C.M., and S.M.B.; Data curation, S.S.P., S.J.R., K.L.M., G.L.S., M.L., F.J., C.K., C.S., G.S., C.M., and S.M.B.; Writing – original draft, S.S.P. and S.M.B; Writing – Review and Editing, S.S.P., S.J.R., K.L.M., G.L.S., M.L., F.J., and S.M.B.; Visualization, S.S.P. and S.M.B; Supervision, S.J.R., C.M.B., G.S., C.M. and S.M.B.

## METHODS

### Cell Culture and generation of Mouse embryonic fibroblasts (MEFs)

HEK293T cells (ATCC; CRL-3216), HEK293 cells (ATCC; CRL-1573), A549 cells (ATCC; CCL-185), Human hepatoma cell line (Huh7), Vero cells (ATCC; CCL-81), Hela cells (ATCC; CCL-2) and PINK1/Parkin modified Hela cells (Ordureau et al., 2014), Primary Dermal Fibroblast (ATCC; PCS-201-012) and murine embryonic fibroblasts (WT MEF), Ajuba^−/−^ and PINK1^−/−^ MEFs were grown in Dulbecco’s modified enrichment medium (GIBCO; 11995-065) containing 10% fetal bovine serum (GIBCO; 16000-044) and 1% penicillin/streptomycin (GIBCO; 15140-122) in an atmosphere of 5% CO2 at 37°C. For WT, PINK1^−/−^ and Ajuba^−/−^ MEFs isolation, fifteen-day old mouse embryos were collected, their separated torsos were washed with PBS, minced, placed in 0.25 % trypsin-EDTA (Invitrogen) containing 1 μg/ml DNase I (Ambion) and incubated at 37°C for 15 min. Cells were filtered using a 100 μM nylon strainer, centrifuged (500 *x g*, 5 min), resuspended in complete DMEM, plated in tissue culture flasks with designation of passage 1. These cells were used for experiments between passage 2-5. All cells were counted on an Image Cytometer (Invitrogen). Cells were either mock infected or infected with SENV (200 Units/well; Charles River Laboratories), VSV (MOI 0.01) and ZIKV PRABC59 strain (MOI 1). Virus was allowed to adsorb for 1 hour at 37°C followed by a media change and incubation for the indicated time.

### Plasmids, siRNAs and Transfections

Human Ajuba and LIMD1 genes were PCR amplified from cDNA templates and directionally cloned into the Gateway entry vector pENTR/SD/D-TOPO (ThermoFisher). Mammalian expression plasmids were then obtained by recombination into Gateway destination vectors pcDNA6.2/FLAG-DEST, GFP-DEST and RFP-DEST (for N-terminal FLAG tag, N’-GFP tag and N’-RFP tag). All plasmids were verified by DNA sequencing. Additional plasmids used to express GFP, MAVS and HA-Ubiquitin were obtained from Addgene (#135046, #52135 and #17608) while EGFP-Parkin and its mutant was a kind gift from Dr. Wolfdieter Springer (Mayo Clinic, Jacksonville FL). ZIKV virus NS5 expression plasmids are previously described (Grant et al., 2016). For siRNA experiments, A549 cells in a 12-well format were transfected with 10 pmol of siRNA (Dharmacon; SMART pool against AJUBA, LIMD1 mRNA and a nonspecific control sequence) for 48 h using Lipofectamine RNAiMAX (Life Technologies). Plasmid transfections were performed according to manufacturer’s protocol in 12 -well plates or 4-well Lab Tek II chamber slides (Nunc) using jetPRIME reagent (VWR INTERNATIONAL; # 89129-924). For harder to transfect Huh7 cells a spinning method was performed post transfection (Spinfection) that adds centrifugation of transfected cells at 1000 rpm for 30 minutes at rt.

### Western Blotting

Cells were washed in phosphate buffered saline (PBS) and harvested in 300 μl each (for 12 well plate) and 500 μl each (for 6 well plate) in RIPA buffer (50 mM Tris-HCl, 150 mM NaCl, 0.1 % SDS, 1% NP-40, 0.5% Na-deoxycholate and DNase I) with complete protease and phosphatase inhibitor cocktail (Roche). Cellular debris was removed by centrifugation (10000 *x g* for 10 min at 4°C) and the supernatant was mixed with sample buffer (2X SB, 62.5 mM TRIS pH 6.8, 10% glycerol, 15 mM EDTA, 4% 2-ME, 2% SDS, and bromophenol blue) and incubated for 10 min at 95°C. An equal amount of sample was resolved by electrophoresis in the presence of SDS on polyacrylamide gels (ThermoFisher). Proteins were transferred to nitrocellulose/PVDF membrane using the iBlot Gel Transfer Device (ThermoFisher) or wet transferred using 0.5 M sodium phosphate transfer buffer (at a constant 1 Amp for an hour using a Bio-Rad apparatus). Membranes were blocked in 5% milk in PBS-T and probed with primary antibody (overnight at 4°C) followed by an hour incubation with secondary antibody at rt with 3X washes of PBST for 10 minutes after each step of incubation. Membranes were blocked in membrane blocking solution (ThermoFisher) with 50mM NaF added when phospho-specific antibodies were used. Immunoreactive proteins were detected by the ECL Plus chemiluminescent system (Thermo Fisher). Western blots were scanned using FluorChem E system (Protein Simple) and quantification of immunoblot bands was performed using ImageJ software.

### Antibodies

The following primary antibodies were used: mouse anti FLAG (#F1804-200UG, Sigma), rabbit anti FLAG tag (#8146S and #14793S, Cell Signaling Technology), mouse anti HA tag (#901502, Biolegend), rabbit anti HA tag (#3724S, Cell Signaling Technology), mouse anti GFP (#632381, Takara Bio Clontech), mouse anti Vinculin (#V9131-100UL, MilliporeSigma), mouse anti GAPDH (#sc-47724, Santa Cruz), mouse anti Ajuba (#sc-374610, Santa Cruz), rabbit anti Ajuba (#34648S and #4897S, Cell Signaling Technology), rabbit anti p-Ubiquitin Ser65 (#ABS1513-I, EMD Millipore), rabbit anti p-Ubiquitin Ser65 (#37642S, Cell Signaling Technology), mouse anti Mitofusin-2 (#sc-100560, Santa Cruz), rabbit anti Mitofusin-1 (#sc-50330, Santa Cruz), rabbit anti Mitofusin-1 (#14739S, Cell Signaling Technology), mouse anti MAVS (#ENZ-ABS259-0100, Enzo), rabbit anti TIM44 (#ab24466, Abcam), rabbit anti-TOMM20 (#HPA011562, Sigma), rabbit anti Calreticulin (#JM-3077-100, MBL international Corporation), rabbit anti PINK1 (#ab23707, Abcam), mouse anti PINK1 (#sc-517353, Santa Cruz), sheep anti p-PINK1 Thr-257 (#68-0057-100, Ubiquigent), mouse anti Parkin (#sc-32282, Santa Cruz), rabbit anti Parkin (#2132S, Cell Signaling Technology), rabbit anti PKR (#ab184257, Abcam), rabbit anti p-PKR Thr-451 (#07-886, EMD-Millipore), rabbit anti eIF2a (#9722S, Cell Signaling Technology), rabbit anti p-eIF2a S51 (#9721S, Cell Signaling Technology), rabbit anti ATF4 (#11815S, Cell Signaling Technology), dsRNA antibody J2 (#10010200, English& Scientific Consulting), mouse anti ZIKV Envelope (#BF-1176-56, BioFront Technologies) and chicken antibody to SENV (#ab33988, Abcam). Mouse isotype IgG control (#sc-2025, Santa Cruz) and rabbit isotype IgG control (#sc-2027, Santa Cruz) was used as a control for immunoprecipitation. Following secondary antibodies were used: HRP conjugated goat anti-mouse antibody (#P0447, Dako from Agilent), HRP conjugated goat anti-rabbit antibody (#P0448, Dako from Agilent), anti-chicken antibodies (#12-341, Millipore) and anti-sheep HRP Secondary antibody (#ab97125, Abcam).

### Inhibitors

Cell culture grade proteasomal inhibitors epoxomicin (Sigma; #E3652) and lysosomal inhibitor bafilomycin A1 (Baf-A1) (Sigma; #B1793) was used at concentration of 200nM each. Oxidative stress inducers tunicamycin (Sigma; # SML1287) and oligomycin A (Sigma; # 75351) were used at 5μg/ml for 5 h in incomplete media. Mitochondrial depolarizing agent carbonyl cyanide 3-chlorophenylhydrazone, CCCP (Millipore Sigma; #C2759) was used at 10μM. Integrated stress response inhibitors ISRIB (Sigma; #SML0843), PKR inhibitor C16 (Millipore Sigma; #527450) and a PKR Inhibitor negative control (Millipore Sigma; #527455) were used at 500nM for 24 h prior to the time of harvest. Ethidium bromide solution (Sigma; #E1385) was used at 100ng/ml.

### Generation of Knockout Mice

Ajuba knockout mice (*Ajuba^−/−^*) were generated using CRISPR/Cas9 technology. Briefly, in vitro synthesized guide RNA (sgRNAs) that were designed to cut shortly after the translation initiation codon in Ajuba Exon 1 were ordered from ThermoFisher’s sgRNA service (<u>CCG</u>GAGTCCGAGAGTCTCAACTT). sgRNA (20ng/ul) was microinjected with Cas9 mRNA (50ng/ug purchased from TriLink BioTechnologies) into the cytoplasm of fertilized eggs collected from C57BL/6N mice (Charles River Laboratory). The injected embryos were cultured overnight in M16 medium, and those that reached 2-cell stage of development were implanted into the oviducts of pseudopregnant foster mothers (CD-1 mice from Charles River Laboratory). Offspring born to the foster mothers were genotyped by PCR and sanger sequencing. Founder mice with desired mutations were bred with C57BL/6NJ mice to establish the knockout mouse line. The two lines #1 (with 4 nt deletion) and #2 (with 22 nt deletion in exon 1) were designated as Ajuba^−/−^ 8216A and Ajuba^−/−^ 8258B respectively. The PINK1 knockout mice (PINK1^−/−^) were procured from Jackson Laboratory.

### Mouse experiments

All animal experiments were approved by the IACUC of Rocky Mountain Laboratories, National Institutes of Health and carried out by a certified staff in an Association for Assessment and Accreditation of Laboratory Animal Care International accredited facility, according to the institution’s guidelines for animal use, and followed the guidelines and basic principles in the U.S. Public Health Service Policy on Humane Care and Use of Laboratory Animals and the Guide for the Care and Use of Laboratory Animals. Male and female mice (4-5 weeks old) were treated by intraperitoneal injection with 2 mg of an anti-mouse IFNAR1 blocking antibody (MAR1-5A3, from Leinco Technologies). The next day, mice were inoculated subcutaneously (via footpad) with 10^3^ PFU of mouse-adapted ZIKV-Dak-41525 (Gorman et al., 2018) kindly provided by Dr. Michael Diamond (University of Washington School of Medicine in Saint Louis). Tissue and blood samples were collected on day 3 and 5 post infection.

### RNA Isolation and quantitative RT-PCR

Total RNA was isolated from cells using RNeasy kit with genomic DNA elimination (QIAGEN). RNA was reverse transcribed using a SuperScript VILO cDNA synthesis kit (ThermoFisher) according to manufacturer’s protocol. cDNA was then used as a template in TaqMan-PCR reactions per manufacturer’s instructions (Applied Biosystems) to quantify mRNA specific for IFNβ (assay ID: Hs01077958_s1), housekeeping gene HPRT (assay ID: Hs01003267_m1), Ajuba (assay ID: Hs00262750_m1) and LIMD1 (assay ID: Hs01040528_m1). Reactions for Real-time RT-PCR were set up in triplicate, cycled and data was collected on the Applied Biosystems GeneAmp 9500 Sequence detection system. Gene expression was normalized to HPRT mRNA levels and expressed as fold change relative to RNA samples from control cells using the comparative ΔCT method.

### Plaque assay

Viral titers from supernatants collected from cells infected with VSV or ZIKV were determined by plaque assay. Briefly, 24 hr prior to titrations, 24-well plates were seeded with 2 × 10^3^ Vero cells per well. Viral samples were 10-fold serially diluted in completed DMEM ranging from 10^−1^ to 10^− 8^ and 125 μl of dilution was added to individual wells. After the plates were incubated for 1h of virus adsorption, the inoculum was removed, and the cells were overlaid with Minimum Essential Medium containing 1.5% carboxymethylcellulose (w/v). The plates were incubated at 37°C for 4 days and were then fixed with 10% formaldehyde for 1 hr at rt followed by staining with 1% crystal violet (in 25% ethanol) for 10 min. Excessive crystal violet and residual overlay media was washed with water and visible plaques were counted to calculate viral titers as plaque forming units per ml (PFU/ml). For analysis of viral distribution in tissues, mice were euthanized at 3 and 5 dpi, and indicated tissues were collected. Organs were individually weighed, homogenized, and prepared as 10% (w/v) suspensions in DMEM/2% FBS/Pen/Strep. Suspensions were then clarified by centrifugation (4,000 rpm for 5 min at 4°C), and the supernatants were titrated using plaque assay.

### Confocal Microscopy

80,000 cells were seeded onto each well of 4 well Lab-Tek II chamber slides (Thermo Fisher Scientific) overnight. To fix, cells were washed with PBS and subsequently fixed with 4% paraformaldehyde for 10 min. Cells were permeabilized with 0.1% Triton X-100 for 5 min at RT, and incubated with blocking solution (PBS, 0.5% BSA, 1% goat serum) for an additional 30 min. Cells were then incubated with primary antibody overnight at 4°C (or 2 hours at RT), washed three times with PBS and further incubated with secondary antibody (in blocking buffer) conjugated to Alexa 488, 594 or 647 (Thermo Fisher Scientific) for 1 h. Slides were washed three times with PBS and once with miliQ water, and mounted onto glass coverslips using Prolong Gold Antifade Reagent with DAPI (Invitrogen). Processed slides were imaged using a Zeiss LSM710 confocal microscope and further analyzed using Zen software (Carl Zeiss).

### Mitochondrial Fractionation

Mitochondria Isolation Kit for Cultured Cells (Thermo Fisher Scientific; # 89874) was used for mitochondrial fractionation using the manufacturer’s protocol. In short, around 2 × 10^7^ cells were pelleted by centrifugation in a 2.0mL microcentrifuge tube at 850 *× g* for 2 minutes. Supernatant was discarded and 800 μL of Mitochondria Isolation Reagent A (with proteasomal inhibitor) was added. Cells were vortexed at medium speed for 5 seconds and incubated on ice for exactly 2 minutes. 10 μL of Mitochondria Isolation Reagent B was added and sample was vortexed at maximum speed for 5 seconds followed by an incubation on ice for 5 min with an in between vortexing step at maximum speed every minute. 800 μL of Mitochondrial Isolation Reagent C (with proteasomal inhibitor) was added and the sample tube was inverted several time to mix properly. The sample was centrifuged at 700 *×g* for 10 minutes at 4°C and the supernatant was transferred to a new 2.0 ml tube, followed by another centrifugation step at 12,000 *×g* for 15 min at 4°C. The pellet contains the isolated mitochondria and supernatant is cytosolic fraction. Mitochondrial pellet was given an additional wash using 500 μL of Mitochondria Isolation Reagent C and centrifugation step at 12,000 *×g* for 5 minutes. The pellet was dissolved directly in 60 μL of lamelli buffer for western blotting as a downstream application and meanwhile stored in −20°C.

### Extracellular Flux (Seahorse) Analysis

MEF isolated from WT, PINK1^−/−^, or and AJUBA^−/−^ mice were seeded at 2×10^4^ cells per well in a XFe96 tissue culture plate and incubated for 24 hours in cDMEM. Cells were washed 2 times with 200 μL of extracellular flux assay medium (DMEM with 25 mM glucose, 2 mM sodium pyruvate, and 2 mM L-glutamine for mitochondrial stress test (Agilent Technologies). Assay medium was then added to each well to make the final well volume 180 uL. Cells were incubated for 1 hr at 37℃ in a non-CO2 incubator prior to extracellular flux analysis. Oxygen consumption rate (OCR) rate was measured using the Mito Stress Test according to manufactures instructions. Briefly, mitochondrial stress assessment included analysis of basal OCR and OCR following injection of oligomycin (2 μM, MilliporeSigma), fluoro-carbonyl cyanide phenylhydrazone (FCCP; 2 μM; Cayman Chemical), and rotenone/antimycin (0.5 μM final concentration for both; MilliporeSigma). All extracellular flux assays were performed on the Seahorse XFe96 Analyzer (Agilent Technologies).

### Flow cytometry

Cells were harvested after 16 hours of treatment conditions, and 30 minutes of mitotracker staining (200nM)(Sprung et al., 2018) followed by staining with LIVE/DEAD Fixable Aqua Dead Cell Stain Kit (ThermoFisher). Data was acquired on a LSRII flow cytometer (BD Biosciences) and analyzed using FlowJo software and are representative of four independent experiments performed. Dead cells, debris and doublets were excluded from all analyses.

### Protein Purification (By GeneScript)

Ajuba and PINK1 DNA sequence was codon optimized and synthesized to be cloned into pET30a vector with N terminal His tag for protein expression in *E. coli*. Transformed *E. coli* strain BL21(DE3) was inoculated into TB medium containing kanamycin and cultured at 37 °C. When the OD600 reached about 1.2, cell culture was induced with IPTG at 15°C for 16 h. Cells were harvested by centrifugation. Cell pellets were resuspended with lysis buffer followed by sonication. The inclusion bodies after centrifugation were dissolved using urea. Target protein from denatured supernatant were refolded and sterilized by 0.22μm filter. Protein concentration was determined by Bradford protein assay with BSA as standard. The protein purity and molecular weight were determined by standard SDS-PAGE along with western blot confirmation.

### Co-immunoprecipitation assay (co-IP)

Cells were washed with PBS and lysed in RIPA buffer (supplemented with 5μg/ml DNAse I and protease inhibitor cocktail). Samples were subjected to centrifugation for 10 min to remove cellular debris and 100 μL of supernatant was reserved for input analysis. Cell lysates were then pre-cleared by addition of Dynabeads Protein G (ThermoFisher) and rotated at 4°C for 3 h. Beads were removed by centrifugation, and 2 μg antibody (corresponding to the protein of interest/ protein tag) was added to each lysate for 2 h with rotation at 4°C. 50 μL of Dynabeads Protein G were added again, and lysates were incubated with rotation at 4°C overnight. Lysates were discarded after a brief centrifugation, and beads were washed 3 times in RIPA buffer for 15 min (at 4°C with rotation) prior to elution by incubation at 95°C for 10 minutes in 50 μL of 2X sample buffer. The eluted samples were assayed by immunoblotting as described above. IP with *E.coli* purified PINK1 and Ajuba used 250nM of each protein in PBS followed by the regular protocol past the preclear step.

### *In-vitro* PINK1 phosphorylation assay

Kinase assay was performed using recombinant proteins purified from *E.coli* and commercially available 10X Kinase buffer (Cell Signaling Technology, #9802). Recombinant proteins (N’-His PINK1 and N’-His Ajuba) were thawed on ice. In a clean microtube, 500 μM ATP in kinase assay buffer diluted in miliQ water to a final concentration of 1× and PINK1 (300 nM) was mixed with Ajuba, the concentration of which was adjusted by dilution from 50 nM to 300 nM for each separate kinase reaction in 25 μl total reaction volume. All reaction mixtures were normalized with BSA to have 1ug total protein in each condition. Negative controls were prepared using as above with either minus Ajuba or minus PINK1. Reactions were spin down for 10 sec at 1,000 *× g* at 4 °C and then were incubated in heated shaker at 30 °C for 30 minutes. The reaction was stopped by boiling for 5 min at 95 °C in 6× SDS loading buffer and were separated by SDS-PAGE electrophoresis (16% gel). Detection of PINK1 auto phosphorylation was assessed using phospho-specific antibody on a western blot.

### Bio-plex Cytokine Analysis

Sera were collected from mock- and ZIKV-infected mice at 3 and 5 dpi. Primary human dermal fibroblast culture supernatants were collected at the indicated time points after ZIKV infection and Control/C16/ISRIB treatment. Cytokine concentrations in serum were measured using a mouse 23-cytokine Bio-Plex Pro Assay according to the manufacturer’s instructions (BioRad). Concentrations for human cytokines were determined using the ProcartaPlex Multiplex Immunoassay (Thermo Fisher Scientific). Briefly, samples were incubated with magnetic beads coupled to cytokine-specific antibodies for 2 h at rt. Beads were washed, incubated with biotin-labeled secondary antibodies for 30 min, washed again and then incubated with a streptavidin reporter for 30 min. After another round of washing, resuspended beads were read by Luminex 200 Bio-Plex Array Reader (Bio-Rad) to acquire data (Internal bead fluorescence, indicative of each distinct cytokine, and fluorescence intensity of signal).

### Electron Microscopy

Cells were grown on Thermanox coverslips and fixed with 2.5% glutaraldehyde in 0.1 M sodium cacodylate buffer. Samples were processed in a PELCO BioWave laboratory microwave (Ted Pella, Reading, California) by post-fixation with 0.5% osmium tetroxide + 0.8% potassium ferrocyanide in 0.1 M sodium cacodylate buffer, buffer rinse, 1% aqueous tannic acid, water rinse, 1% aqueous samarium acetate, water rinse, dehydration into ethanol, embedment into epon, and polymerization overnight at 60°C. Thin sections were cut with a Leica UC6 ultramicrotome (Leica Microsystems, Vienna, Austria), stained with uranyl acetate, and imaged on either a 120 kV HT7800 (Hitachi) operating at 80 kV with an XR-81B CMOS digital camera (AMT Imaging System, Woburn, Massachusettes) or on a 120 kV Tecnai Bio Twin Spirit (FEI, Hillsboro, Oregon) with a Rio CMOS digital camera (Gatan, Pleasanton, California).

### NGS Library Preparation

Cells were harvested in Trizol and frozen on dry ice. Each sample lysate was combined with additional Trizol to bring the final volume to 1000μl in each sample, 200μl of 1-Bromo-3-chloropropane (MilliporeSigma) was added, samples mixed, and centrifuged at 16,000 × *g* for 15 min at 4°C. RNA containing aqueous phase of 600μl was collected from each sample and passed through Qiashredder column (Qiagen) at 21,000 *x g* for 2 min to homogenize any remaining genomic DNA in the aqueous phase. Aqueous phase was combined with 600μL of RLT lysis buffer (Qiagen) with 1% β mercaptoethanol and RNA was extracted using Qiagen AllPrep DNA/RNA 96-well system. An additional on-column Dnase I treatment was performed during RNA extraction. Standard RNeasy extraction protocol resulted in RNAs larger than 200nt. All sample processing was performed using amplicon-free reagents and tools in aerosol resistant vials. RNA was quantitated by spectrophotometry and RNA yield ranged from 0.6 to 8.5μg. RNA quality was analyzed using Agilent 2100 Bioanalyzer (Agilent Technologies) and RNA integrity number (RIN) ranged from 7.6 to 9.9 (showing an overall exceptional RNA quality). 500 ng RNA was used as input for the TruSeq Stranded mRNA-Seq Sample Preparation Kit (Illumina). The protocol was followed without modification. Final library size distribution was assessed on a BioAnalyzer DNA 1000 chip (Agilent Technologies). The average size of the libraries was on target at around 310 bp. Libraries were quantified using the Kapa SYBR FAST Universal qPCR kit for Illumina sequencing on the CFX384 Real-Time PCR Detection System (Bio-Rad Laboratories). The libraries were diluted to 4 nM stocks and pooled equitably for sequencing. The 4 nM pool of libraries was prepared for sequencing by denaturing and diluting to a 1.8 pM stock for clustering to the flow cell. On-board cluster generation and paired-end sequencing was completed on the NextSeq 550 (Ilumina) using a High Output 150 cycle kit (Illumina). The average cluster density was 177 k/mm^2^ resulting in 420 million reads passing filter per run, with an average of 22.6 million reads per sample. Raw fastq reads were trimmed of Illumina adapter sequences using cutadapt version 1.12 and then trimmed and filtered for quality using the FASTX-Toolkit (Hannon Lab). Remaining reads were aligned to the mouse genome assembly mm10 using Hisat2. Reads mapping to genes were counted using htseq-count.

### Bioinformatic analysis

Briefly, differential expression analysis in heatmaps was performed using the Bioconductor package DESeq2. Cytokine analysis for RNA seq data involves 157 genes mapped on the KEGG pathway mmu04060. Volcano plots were created to identify cytokine genes showing more than 2 log2 fold change with corresponding significant P values in control versus infected samples. Functional and network analysis of statistically significant gene expression changes was performed using Ingenuity Pathways Analysis (IPA, QIAGEN). Significant canonical pathways are identified from IPA based on enrichment of the zscore.

### Mass spectrometry (By MS Bioworks)

Samples were processed by SDS-PAGE using a 4-12% Bis-Tris NuPAGE gel (Invitrogen) with the MOPS buffer system. The target bands were excised and processed by in-gel digestion using a robot (ProGest, DigiLab). The digest was analyzed by nano LC-MS/MS with a Waters NanoAcquity HPLC system interfaced to a ThermoFisher Q Exactive. Peptides were loaded on a trapping column and eluted over a 75μm analytical column at 350nL/min; both columns were packed with Luna C18 resin (Phenomenex). The mass spectrometer was operated in data-dependent mode, with the Orbitrap operating at 60,000 FWHM and 17,500 FWHM for MS and MS/MS respectively. The fifteen most abundant ions were selected for MS/MS. Samples were analyzed in analytical duplicate. Data were searched using a local copy of Mascot (Matrix Science) and Mascot DAT files were parsed into Scaffold (Proteome Software) for validation, filtering and to create a non-redundant list per sample. Data were filtered using a minimum protein value of 99.9%, a minimum peptide value of 50% (Prophet scores) and requiring at least two unique peptides per protein. Scaffold results were exported as mzidentML and imported in to Scaffold PTM in order to assign site localization probabilities using A-score with minimum localization probability filter of 50%.

### RNA scope in-situ Hybridization

In situ hybridization was performed using the RNAScope 2.5 VS assay (Advanced Cell Diagnostics, Newark, CA) according to the manufacturer’s instructions using a mouse Ajuba specific probe (cat#579019).

### Statistical Analysis

All data were evaluated for significance using student T-tests, Mann Whitney test, or one-way/two-way ANOVA with appropriate multiple comparison post tests using GraphPad Prism 7 software. Statistical significance was assigned as when *p* values were <0.05. Specifically, \**p*<0.05, \*\**p*<0.01, \*\*\**p*<0.001, \*\*\*\**p*<0.0001, not significant (NS) when *p* > 0.5.

## Supplementary Figures and Legends

**Supplemental Figure 1:**
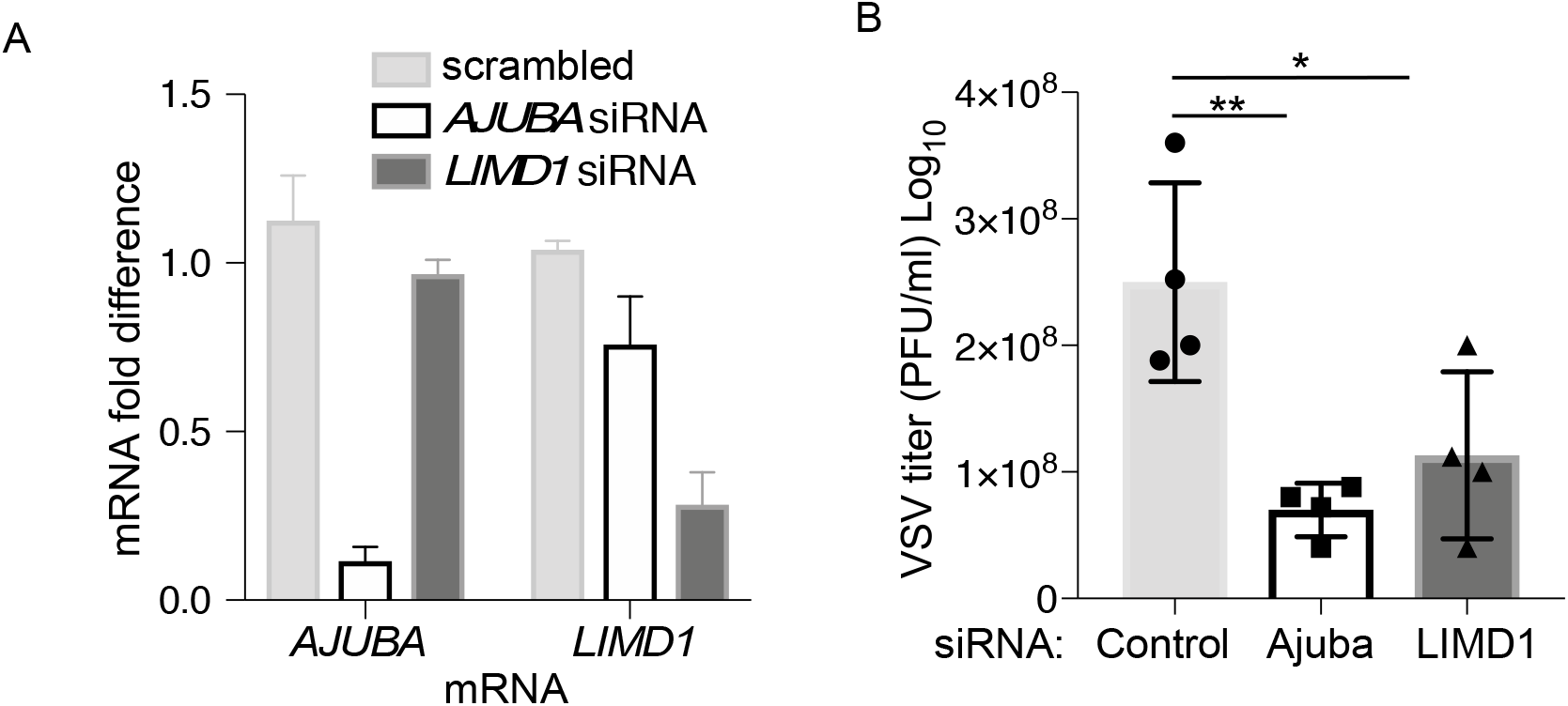
Ajuba negatively regulates MAVS expression. **A.** Efficiency of mRNA depletion of *AJUBA* or *LIMD1* in A549 cells at 48 h post transfection with siRNAs targeting each gene. **B.** Titer of VSV at 24 hpi of cells from C. Error bars represent mean±SD from 2 independent experiments performed in duplicate; *P<0.05, **P<0.01, ***P<0.001 by one-way ANOVA.

**Supplemental Figure 2:**
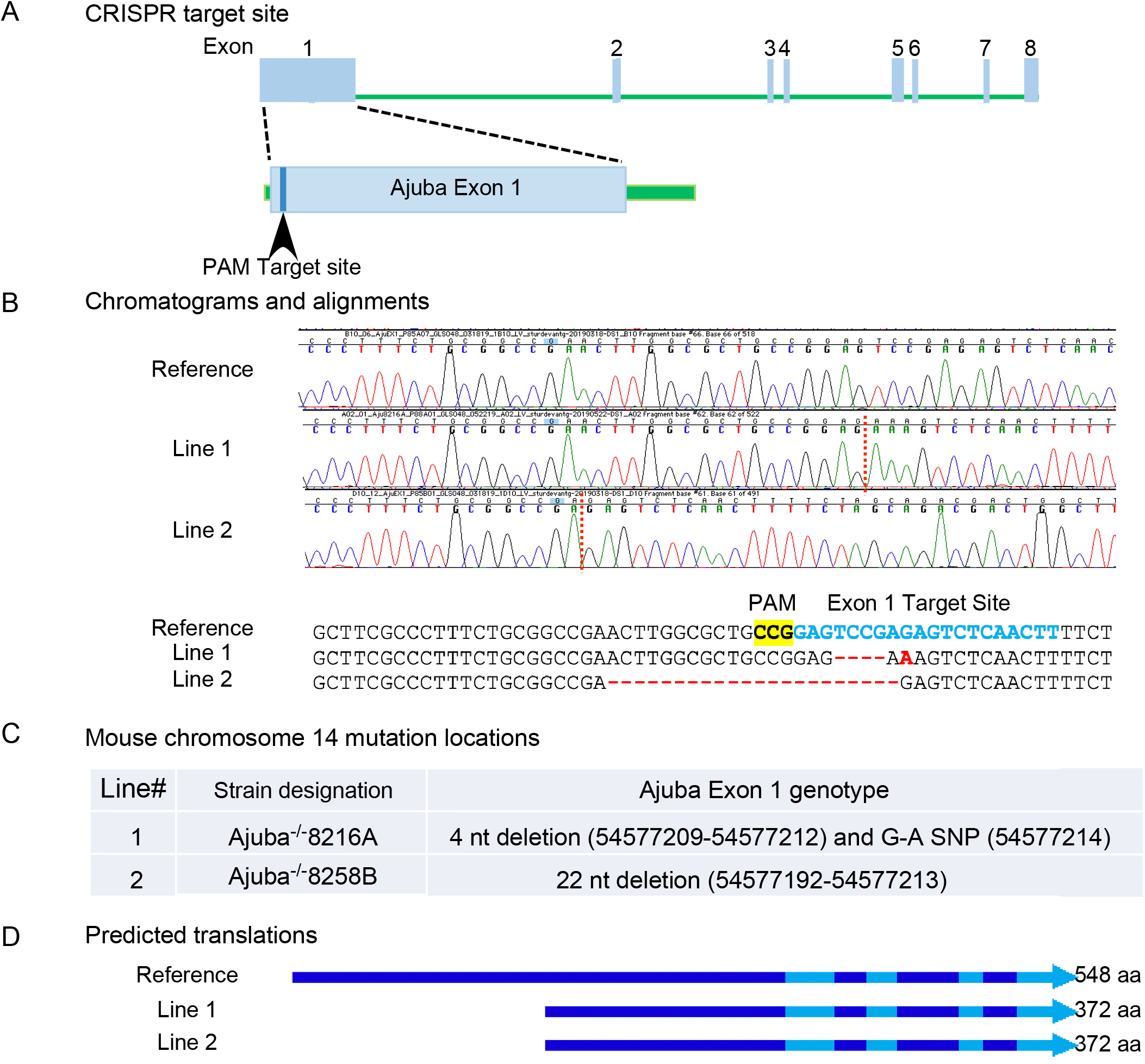
Design and verification of Ajuba^−/−^ mice using CRISPR/Cas9-mediated gene editing. **A.** Exon structure of the *Ajuba* gene. *Ajuba* Exon 1 was targeted by the gRNA sequence CCGGAGTCCGAGAGTCTCAACTT. **B.** Sequence verification of gene disruption in two lines of mice designated 8216A and 8258B. **C.** Summary of sequence modifications and **D.** Predicted potential ORF expression.

**Supplemental Figure 3.**
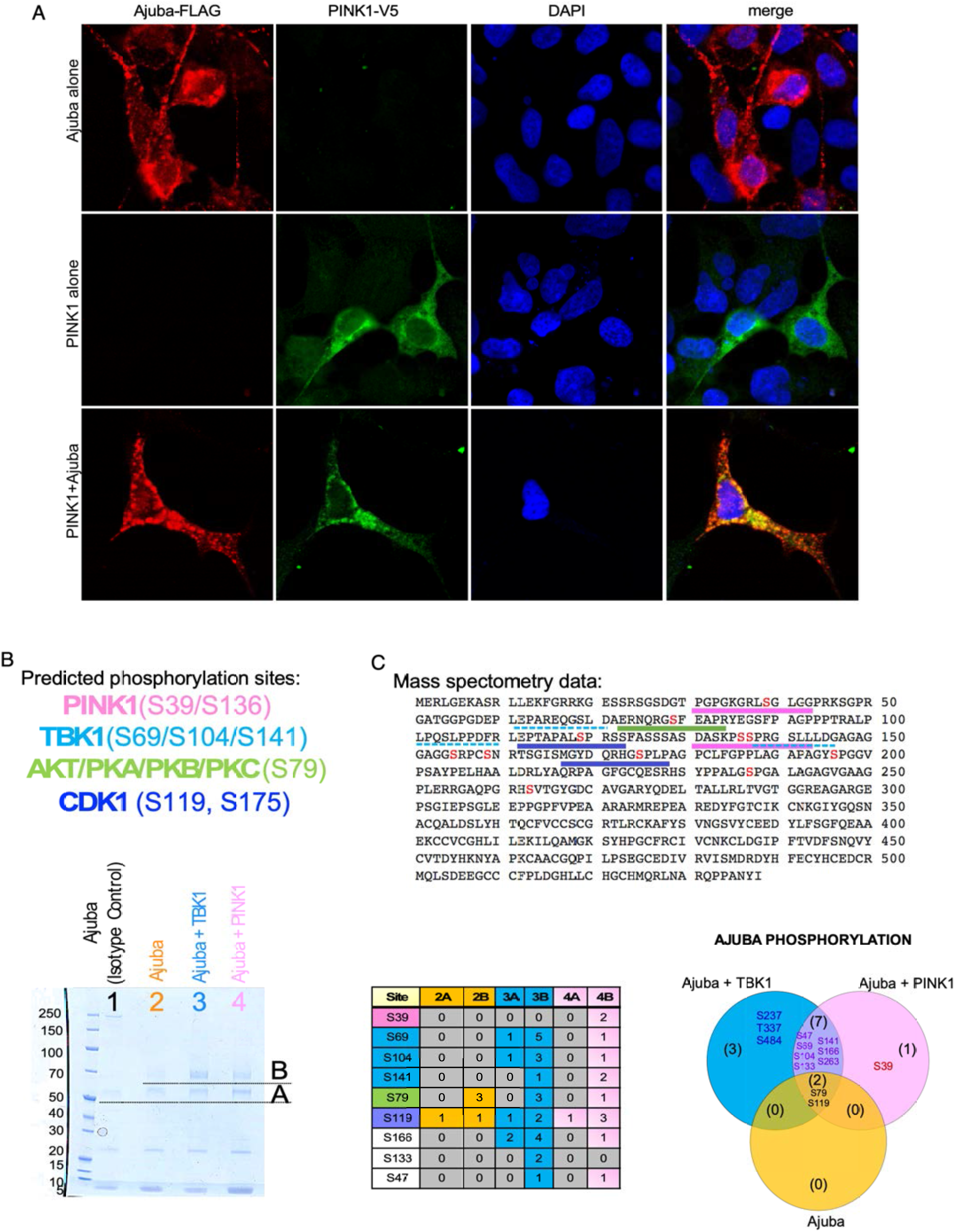
Ajuba is phosphorylated by PINK1. **A.** Confocal microscopy of HEK293T cells expressing FLAG-Ajuba (red), PINK1-V5 (green) or both showing co-localization of the two proteins (yellow). Nuclei were counterstained with DAPI (blue). **B.** Predicted phosphorylation Ser/Thr sites in the pre-LIM region of Ajuba are associated with PINK1, TBK1, AKT/PKA/PKB/PKC, and CDK1. Therefore Ajuba was expressed alone or co-expressed with TBK1 or PINK1 and run on SDS-PAGE. Two bands were excised for analysis by mass spectrometry. The upper band (B) was identified as highly phosphorylated Ajuba. **C.** Ajuba sequence with predicted phosphorylation sites indicated. The table summarizes the mass spectrometry data showing the spectral counts of basally expressed (Band A) and highly phosphorylated Ajuba (Band B). This analysis suggested that Ajuba is basally phosphorylated at S79 and S119. TBK1 induced further phosphorylation at S47, S69, S104, S141, S133 and S166. PINK1 induced specific phosphorylation of Ajuba at S39, but also induced phosphorylation of the TBK1-dependent residues as summarized in the Venn diagram.

**Supplemental Figure 4.**
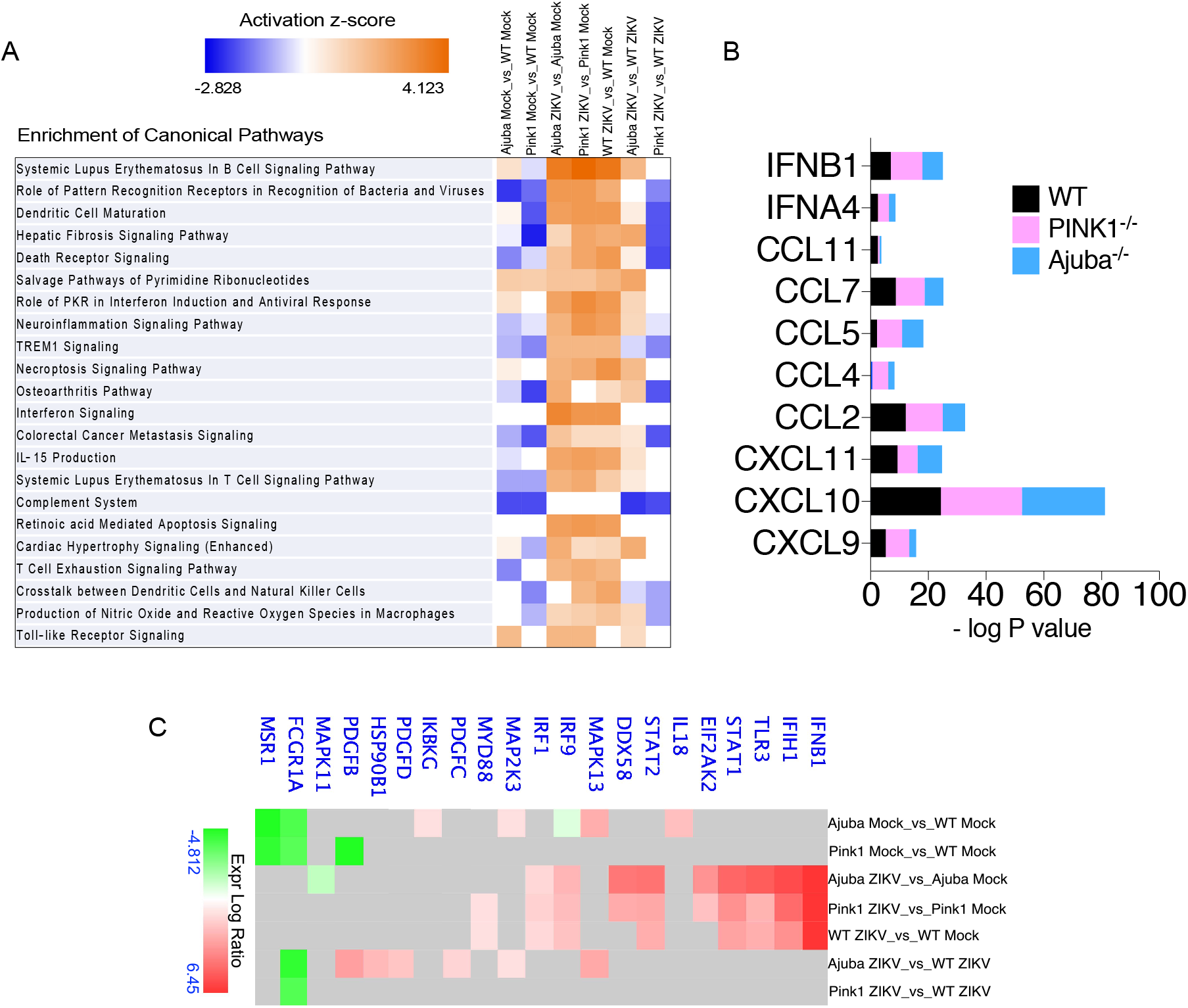
RNAseq and DEG expression in WT, PINK1^−/−^ and Ajuba ^−/−^ MEFs. **A.** Gene Ontology (GO) terms from IPA analysis demonstrating higher expression of interferon and inflammatory pathways in the absence of PINK1 or Ajuba, including the role of PKR in interferon induction and antiviral response. **B.** Log P value of chemokine expression shown in Figure 6E. **C.** Individual genes from the pathway designated as ‘role of PKR (EIF2AK2) in interferon induction and antiviral response’ demonstrating high expression in Ajuba^−/−^ and PINK1^−/−^ MEFs.

**Supplemental Figure 5.**
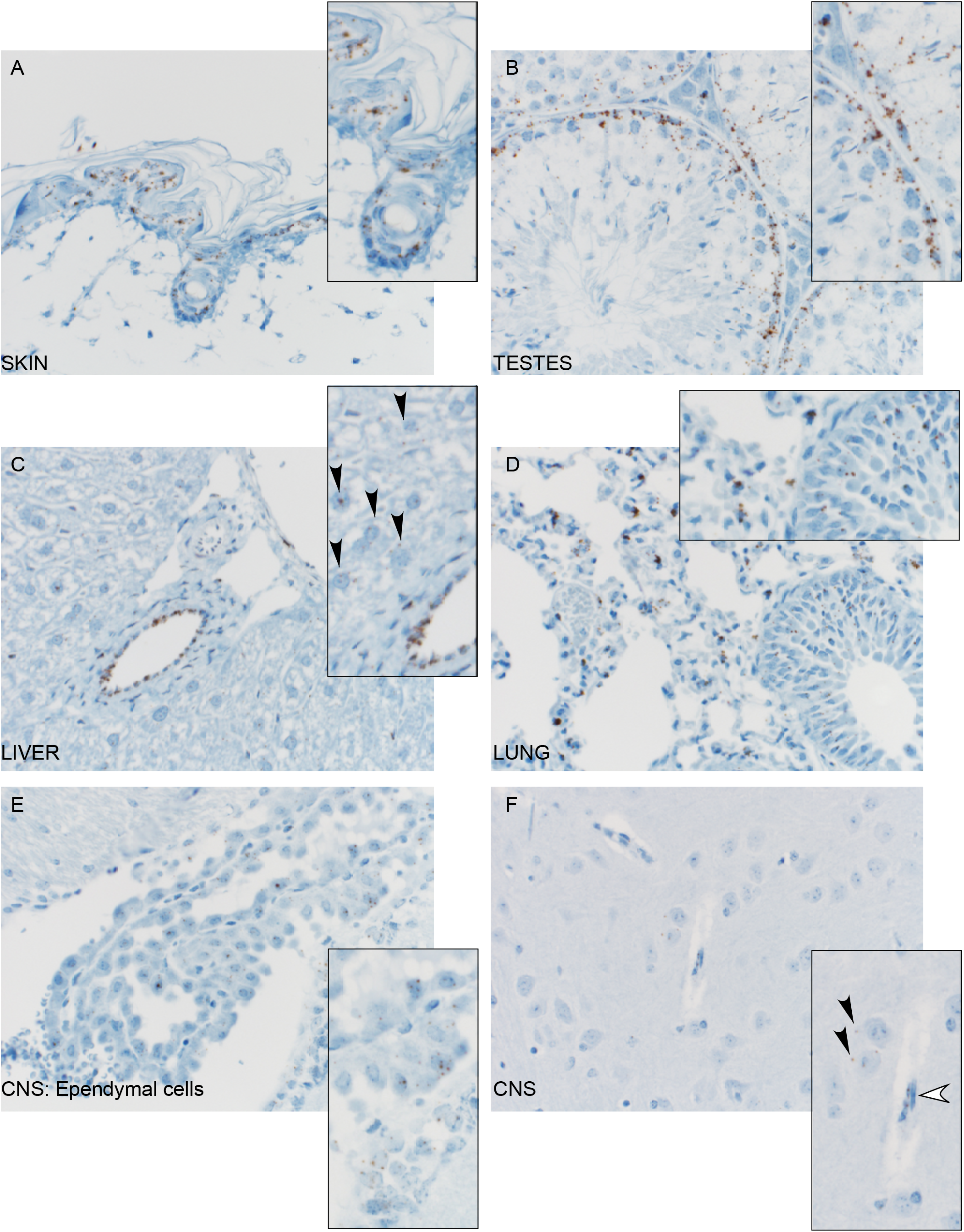
*Ajuba* mRNA expression in mouse tissues by RNAScope. *Ajuba* mRNA staining (brown) was observed in **A.** keratinocytes and endothelial cells in the skin, **B.** Sertoli cells in the testes, **C.** hepatocytes (arrowheads) and endothelial cells in the liver, **D.** epithelial and endothelial cells in the lung, **E.** ependymal cells and **F.** neurons (black arrowheads) and endothelial cells (white arrowheads) in the CNS. Tissue sections were counterstained with hematoxylin.

**Supplemental Figure 6.**
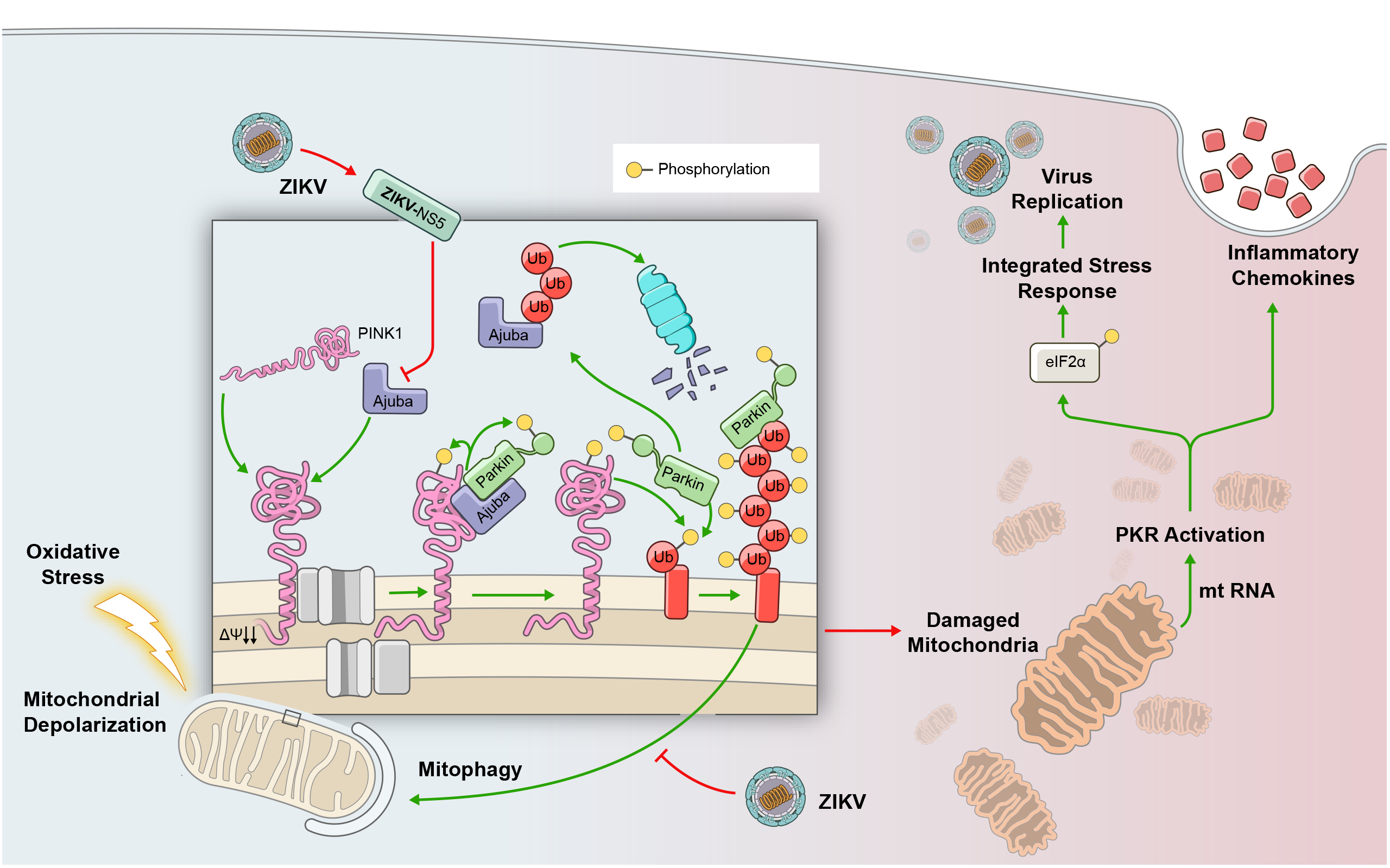
Schematic representation of findings. ZIKV virus suppresses PINK1-Parkin-dendent mitophagy through the actions of NS5 binding to and inhibiting mitochondrial recruitment of Ajuba. This results in mitochondrial RNA release and PKR activation and activation of the ISR to create a cellular environment that favors virus replication. However, PKR-dependent amplification of chemokines and specific cytokines central to the pro-inflammatory response to ZIKV is uncoupled from virus replication.

